# Pancreatic islets undergo functional and morphological adaptation during development of Barth Syndrome

**DOI:** 10.1101/2024.06.28.601122

**Authors:** Christopher Carlein, Markus D. A. Hoffmann, Andressa G. Amaral, Caroline Bickelmann, Ahmadali Lotfinia, Laurie-Anne de Selliers, Johanne Audoze-Chaud, Selina Wrublewsky, Marcel A. Lauterbach, Karina von der Malsburg, Martin van der Laan, Monika Bozem, Markus Hoth, Patrick Gilon, Magalie A. Ravier, Bruce Morgan, Emmanuel Ampofo, Christoph Maack, Leticia Prates Roma

## Abstract

Barth syndrome is a multisystem genetic disorder caused by mutation in *TAFAZZIN*, a gene that encodes a phospholipid:lysophospholipid transacylase important for cardiolipin remodeling. Barth Syndrome patients suffer from a number of symptoms including early heart failure, fatigue, and systemic metabolic alterations, including hypoglycemia. The endocrine pancreas is central to glucose homeostasis, however, the impact of defective cardiolipin remodeling on pancreatic islet function and the consequences for systemic metabolism is unclear. Surprisingly, in a mouse model with global *TAFAZZIN* knockdown, we observed improved glucose tolerance compared to wildtype littermates. We show that pancreatic islet metabolism and secretory function are robustly maintained through various compensatory mechanisms including increased glucose uptake and increased mitochondrial volume. Transcriptomics analyses revealed increased expression of genes encoding proteins involved in N-acetylglucosamine synthesis and protein *O*-linked N-acetylglucosaminylation. These pathways might provide a molecular mechanism for coupling metabolic changes to mitochondrial volume regulation.

## Introduction

Barth syndrome (BTHS) is a life-threatening, X-linked multisystem disorder characterized by pleiotropic phenotypes including heart failure, growth delay, skeletal myopathy and neutropenia (1). BTHS patients also show changes in whole-body fatty acid, glucose and amino acid metabolism(2). BTHS is caused by mutations in the *TAFAZZIN* (*Taz*) gene, which encodes Tafazzin (Taz1), a mitochondrial phospholipid:diacylglycerol transacylase essential for cardiolipin (CL) remodeling (1). The direct consequence of *Taz* mutation is the accumulation of monolysocardiolipin (MLCL), a precursor of CL, which has been shown to have a lower affinity for the respiratory chain complexes III and IV (3, 4). Changes in CL content or composition, i.e. the identity of the associated fatty acids, correlate with defective formation of mitochondrial supercomplexes, aberrant cristae formation and shape, decreased respiration, decreased ATP production and increased reactive oxygen species (ROS) production (4–7). Other cellular functions, including mitophagy and apoptosis, were also shown to be affected by changes in CL. However, the consequences of defective CL remodeling are tissue-specific and many are still unknown or unclear, with contradictory reports found in the published literature (8–10). For example, increased mitochondrial ROS production has been suggested to have a causal relationship with cellular dysfunction in BTHS, while others have reported no changes and no significant impact in the disease (8, 11–13). We recently showed that mitochondrial calcium uptake is strongly decreased in cardiomyocytes from *Taz*-KD mice due to decreased levels of the mitochondrial calcium uniporter (MCU). Consequently, Krebs cycle activation and mitochondrial respiration during β-adrenergic stimulation is impaired, leading to a lack of inotropic reserve in BTHS cardiomyopathy (7, 8). We did not observe any *in vivo* changes in cardiomyocyte H_2_O_2_ levels, likely due to increased antioxidant defense (8), which may be driven by eIF2α/ATF4-mediated upregulation of one-carbon metabolism and a subsequent increase in glutathione production (14).

Metabolic disorders are associated with absolute changes in fatty acid levels as well as changes in the relative abundance of fatty acids with different chain lengths and saturation. Changes are also seen in the fatty acid profile and abundance of glycerophospholipids, including CL. For example, in a streptozotocin (STZ)-induced diabetic mouse model, CL content was strongly decreased in the myocardium (15). Furthermore, STZ treatment induced considerable CL remodeling, with a strong shift from CL enriched in 18:2 fatty acids to 22:6 fatty acid-enriched CL. Similar changes were also observed in an obese ob/ob mouse model (16). These observations suggest a vital role of CL content and fatty acid composition in maintaining metabolic functions and a possible role in the development of metabolic diseases (16). Consistent with this hypothesis, CL was shown to be important for whole-body energy homeostasis by regulating non-shivering thermogenesis and CL levels were shown to positively correlate with insulin sensitivity (17). Interestingly, *Taz-*KD mice are resistant to diet-induced obesity and are protected against hepatic steatosis (18), again supporting a role of Taz1 and CL in whole-body metabolism. BTHS patients display recurrent hypoglycemia and disrupted fatty acid and amino acid metabolism (19, 20).

Pancreatic islets are key players in maintaining whole-body energetic balance and glucose homeostasis, harboring the cells that secrete insulin (β-cells), glucagon (α-cells), and somatostatin (δ-cells). Recently, *Taz* deficiency was reported to lead to decreased islet insulin secretion and oxygen consumption, an effect that was significant in low glucose concentrations (21). However, this observation does not appear to be consistent with the occurrence of frequent hypoglycemic episodes in human patients and raises questions about the role of pancreatic insulin secretion in BTHS metabolic phenotypes.

In this study, we systematically investigated the impact of *Taz* knockdown in the pancreatic β-cell insulin secretion pathway. Using an in vivo *Taz*-KD mouse model, we showed a surprisingly robust preservation of pancreatic islet function, which in part appears to be a consequence of multiple compensatory mechanisms including increased glucose uptake, increased *O*-linked N-acetylglucosamine pathways, and increased mitochondrial volume. *In vivo*, glucose tolerance is enhanced in *Taz*-KD mice compared to WT littermates, without changes in insulin and glucagon levels, pointing to an important role of the peripheral tissues in glucose handling in BTHS.

## Methods

### Animal models

All animal experiments were approved by the local authorities (animal experiment approval 08/2018 and 19/2019) and in accordance with the Society of Laboratory Animal Science (GV-SOLAS) guidelines, following the 3R principles. Male and female mice were used.

#### shTaz

*Tafazzin* knockdown (*Taz*-KD) mice model was obtained from Jackson Laboratories (B6.Cg-Gt(ROSA)26Sor^tm37(H1/tet0-RNAi:Taz)Arte^/ZkhuJ, stock number: 014648). Doxycycline (doxy) in a concentration of 625 mg of doxy/kg was added to the standard rodent chow (A153D70623, Ssniff, Germany) leading to induction of short hairpin RNA (shRNA)-mediated knockdown of *Taz*, as described previously (22).

#### ShTaz x mito-roGFP2-Orp1

The mito-roGFP2-Orp1 mouse strain (first described in (23), a kind gift of Prof. Dr. Tobias Dick), which globally expresses a H_2_O_2_ sensor (ROSA26/CAG-stop^fl^-mito-roGFP2-Orp1 × CMV-Cre) targeted to the mitochondria matrix, was crossbreed with the *Taz*-KD mice.

### Genotyping

The genotype of the animals was identified by polymerase chain reaction (PCR). Tissue samples were incubated in DNA extraction buffer at 65 °C for 15 min, then vortexed and incubated at 98 °C for 2 min. The extracted DNA was amplified using the according primers listed in supplements. All mouse strains used in this article have the NNT protein and hence, are on a C57BL/6N background.

### Pancreatic islet isolation and culture

Isolation of mouse pancreatic islets was performed as previously described (24). Briefly, mice were anaesthetized with isoflurane and sacrificed via cervical dislocation. Afterwards, the body was opened ventrally, and the ampulla connecting the pancreas and small intestine was clamped. Next, the pancreas was perfused via the pancreatic duct using an ice-cold collagenase solution (Collagenase P, 11213865001, Merck/Sigma Aldrich) with a concentration of 0.63 mg/ml. The perfused pancreas was digested for 20 min in a water bath at 37 °C and washed 3x in Krebs-Henseleit-buffer (KHB), containing 24 mM NaHCO_3_, 120 mM NaCl, 4.8 mM KCl, 1.2 mM MgCl_2_, 2.5 mM CaCl_2_, 5 mM HEPES, 0.2% Bovine serum albumin (BSA), 1% Penicillin/Streptomycin (P/S) and 10 mM glucose. Afterwards, pancreatic islets were hand-picked with a 10 µl pipet under a stereo microscope and separated from the exocrine tissue. The collected pancreatic islets were cultured in RPMI 1640 (ref: 11875093, Gibco^TM^) supplemented with 10% (v/v) Fetal Bovine Serum (FBS) and 1% (v/v) P/S under 5% CO_2_ and 37 °C, until further use. The pancreatic islets were cultured for 1 - 3 days and groups of similar size WT and *Taz*-KD islets were formed before each experiment.

### RNA isolation

Pancreatic islets (groups of around 150 per RNA sample) were lysed in ice-cold TRIzol^TM^ reagent (ref: 15596018, Thermo Fisher) and frozen at -80 °C until further use. RNA extraction was performed according to standard protocol and isolated RNA was either sent for RNA sequencing (RNAseq) to Novogene (Cambrige, UK) or reverse transcribed to cDNA.

### Quantitative RT-PCR

Quantitative RT-PCR was analysed using the CFX96 C1000 touch thermocycler (ref: 785BR06498, Bio-Rad) with Taqman specific assays for *Taz* and GAPDH genes. The Taqman assay IDs are listed in the supplements (Suppl. Table 2).

### Sample preparation for lipidomics analysis

Groups of 600 islets per lipidomics sample were collected by pooling all pancreatic islets from 2 - 3 animals with the same genotype and gender. After culture, islets were homogenized by dispersion in calcium- and magnesium-free PBS with additional sonication and centrifugation (chapter: Homogenization of pancreatic islets). Processed samples were kept at -80 °C or protein content was measured using BCA assay. Samples were further analyzed by Lipotype Lipidomics Dresden, Germany).

### Lipid extraction for mass spectrometry lipidomics

Mass spectrometry (MS)-based lipid analysis was performed by Lipotype Lipidomics GmbH (Dresden, Germany) as described (25). Lipids were extracted using a chloroform/methanol procedure (26). Samples were spiked with internal lipid standard mixture containing: cardiolipin 14:0/14:0/14:0/14:0 (CL), ceramide 18:1;2/17:0 (Cer), diacylglycerol 17:0/17:0 (DAG), hexosylceramide 18:1;2/12:0 (HexCer), lyso-phosphatidate 17:0 (LPA), lyso- phosphatidylcholine 12:0 (LPC), lyso-phosphatidylethanolamine 17:1 (LPE), lyso- phosphatidylglycerol 17:1 (LPG), lyso-phosphatidylinositol 17:1 (LPI), lyso-phosphatidylserine 17:1 (LPS), phosphatidate 17:0/17:0 (PA), phosphatidylcholine 17:0/17:0 (PC), phosphatidylethanolamine 17:0/17:0 (PE), phosphatidylglycerol 17:0/17:0 (PG), phosphatidylinositol 16:0/16:0 (PI), phosphatidylserine 17:0/17:0 (PS), cholesterol ester 16:0 D7 (CE), sphingomyelin 18:1;2/12:0;0 (SM), triacylglycerol 17:0/17:0/17:0 (TAG). After extraction, the organic phase was transferred to an infusion plate and dried in a speed vacuum concentrator. The dry extract was re-suspended in 7.5 mM ammonium formate in chloroform/methanol/propanol (1:2:4; V:V:V). All liquid handling steps were performed using Hamilton Robotics STARlet robotic platform with the Anti Droplet Control feature for organic solvents pipetting.

### MS data acquisition

Samples were analyzed by direct infusion on a QExactive mass spectrometer (Thermo Scientific) equipped with a TriVersa NanoMate ion source (Advion Biosciences). Samples were analyzed in both positive and negative ion modes with a resolution of R_m/z=200_=280000 for MS and R_m/z=200_=17500 for Tandem mass spectrometry (MSMS) experiments, in a single acquisition. MSMS was triggered by an inclusion list encompassing corresponding MS mass ranges scanned in 1 Da increments (27). Both MS and MSMS data were combined to monitor CE, DAG and TAG ions as ammonium adducts; LPC, LPC O-, PC and PC O- as formiate adducts; and CL, LPS, PA, PE, PE O-, PG, PI and PS as deprotonated anions. MS only was used to monitor LPA, LPE, LPE O-, LPG and LPI as deprotonated anions, and Cer, HexCer and SM as formiate adducts.

### Lipidomics data analysis and post-processing

Data were analyzed with an in-house developed lipid identification software based on LipidXplorer (25, 28). Data post-processing and normalization were performed using an in-house developed data management system. Only lipid identifications with a signal-to-noise ratio >5, and a signal intensity 5-fold higher than in corresponding blank samples were considered for further data analysis.

### Glucose tolerance test (GTT)

After a 6 h fasting period, mice were injected with 2.2 mg glucose (glucose monohydrate, ref: 6780.1, Carl Roth) solution per gram of body weight into the peritoneum (IP injection). Plasma glucose levels were monitored throughout the experiment (time points: 0, 7, 15, 30, 60 and 120 min) using a glucosimeter (Accu-Chek^®^, ref: 06870333001, Aviva). In some experiments, at each time point, 15 µl of blood were collected from the tail of the mice in an ethylenediaminetetraacetic acid (EDTA) coated tube (Microvette^®^ CB 300 K2E, ref: 16.444.100, Sarstedt). Using centrifugation, the blood plasma was separated at 2000 rcf for 15 min at 4 °C. Additionally, the samples were centrifuged again at 1000 rcf for 5 min to remove any possible remaining pellet. The supernatant was frozen at -80 °C. Analysis of plasma glucagon and insulin levels were performed with the corresponding mouse plasma insulin (HTRF insulin mouse serum kit, ref: 62IN3PEF, Cisbio/Perkin Elmer) and glucagon (mouse glucagon ELISA kit, ref.: 81518, Crystal Chem) kit.

### Immunohistochemistry

#### Cryoslices

The number of α-, β-, or δ-cells of the *in vivo Taz-*KD model were quantified by immunohistochemistry (IHC). After isolation of the whole pancreas, the tissue was washed in PBS and fixed in 4% Perfluoroalkoxy alkanes (PFA) overnight at RT. On the next day, the tissue was washed for 4 h in PBS, before it was transferred to a 30% sucrose solution and kept for 3 h. Subsequently, the whole pancreas was rapidly frozen in Tissue-Tek^®^ O.C.T.^TM^ (SA62550-01, Science services) using liquid nitrogen-cooled isopentane. Cutting was performed at a Leica cryostat setting a thickness of 5 µm and a temperature of -15 °C. Pancreas slices were dried for 30 min, before being washed with PBS and treated with 3% goat serum for 1 h in a wet chamber. Subsequently, 50 µl of primary anti-insulin together with anti-glucagon or anti-somatostatin antibodies (Suppl. Table 1, 1:200 diluted) were added and incubated overnight in wet chamber at 4 °C. On the following day, slides were again washed with PBS and 50 µl of Alexa 594 and Alexa 488 secondary antibodies (Suppl. Table 1, 1:400 diluted) were added for 1 h 15 min at RT. After a final washing step slides were dried and mounted with Dako mounting media. Imaging was performed with the Axio Observer 7 microscope (Zeiss, Germany) using a 20x air objective.

### Paraffin slices

Paraffin embedding was either performed on the whole pancreas of shTaz animals of the *in vivo* model or on isolated pancreatic islets of the *in vitro* shTaz model. Freshly isolated pancreatic islets were clotted before PFA fixation. First, the pancreatic islets were incubated in 3 ml RPMI 1640 medium on top of 1 ml agarose overnight at 37 °C and 5% CO_2_. Subsequently, the pancreatic islets were clotted in a mixture of human platelet-poor plasma, Hepatoquick and 10% CaCl_2_. The clotted pancreatic islets and the whole pancreas were placed for 24 h in PFA and then embedded with an ethanol, xylol and paraffin protocol in a tissue processor (SLEE medical GmbH, Germany). Paraffin blocks were cut with a microtome and stained against different cell types (namely α-, β-, or δ-cells), Ki67 and cleaved caspase- 3 (Suppl. Table 1). After antibody staining, the slides were treated with DAPI (ref: 10116287, Thermo Fisher) to visualize the nuclei. Imaging was performed with the Axio Observer 7 system (Zeiss, Germany) using a 20x air objective. The analysis was performed in ImageJ and positive stained cells were counted using the cell counter plugin in ImageJ.

### Static insulin and glucagon secretion

Static insulin and glucagon secretion experiments were performed with isolated pancreatic islets after overnight culture. The static experiments were performed in 24-well plates. Initially, all pancreatic islets were pre-incubated under low glucose conditions (2.8 mM) in KHB for 45 min (37 °C, 5% CO_2_). Subsequently, groups of 10 islets were incubated at different glucose concentrations (2.8, 5.6, 10, 20 mM) for 1 h (37 °C and 5% CO_2_). After the incubation, supernatant was collected and frozen at -20 °C until further analysis. The islets that have been incubated in 20 mM glucose KHB were transferred to a pre-warmed 24-well plate with 0.5 mM glucose KHB to stimulate glucagon secretion. After 1 h at 37 °C and 5% CO_2_, supernatant with secreted glucagon was frozen at -20 °C. Subsequently, the islets were dissolved in 74% EtOH, 0.5% HCl and 25.5 % ddH_2_O for extracting insulin content, which were frozen at -20 °C. Analysis of supernatant insulin and glucagon levels was performed using a HTRF insulin ultra- sensitive (ref: 62IN2PEG, revvity) and HTRF glucagon (ref: 62CGLPEG, revvity) kit.

### Dynamic insulin secretion

The dynamic insulin secretion was performed using a homemade microscope chamber. Before the dynamic insulin experiments, the pancreatic islets were starved with 2 mM glucose in KHB and placed into the microscopy chamber. A peristaltic pump system (Ismatec^®^ Reglo ICC Digital peristaltic pump, ref: ISM4412, Cole-Parmer) was set a flow of 1 ml/min and the buffers were kept warm (37 ± 2 °C) with a bead bath (BeadBath Duo, 2 l bath, ref: G20001365, Biozym). With the start of the pump system the low glucose solution (2 mM) was pumped into the chamber. The outflowing liquid was collected in ice-cold tubes. The initial 5 min were pooled and represented the baseline of the dynamic insulin experiments. Afterwards, the outflowing liquid was collected each minute in a new reaction tube. After 10 min, a 20 mM glucose KHB solution was pumped into the chamber. 20 min into the experiment, the outflowing liquid was collected every 2 minutes. After 30 min, a 30 mM KCl solution was applied, and the outflowing liquid was again collected every minute in a new reaction tube. At the end of the experiment, all collected solutions were frozen at -20 °C. Additionally, the pancreatic islets were removed from the microscope chamber and frozen in TE buffer at -20 °C. The insulin secretion levels were assessed using the HTRF insulin ultra-sensitive kit (ref: 62IN2PEG, revvity) and normalized to DNA content using the Pico488 dsDNA quantification kit (NBX-76675, Lumiprobe).

### Dispersion of pancreatic islets

Dispersion of pancreatic islets was performed by transferring the islets (50 islets per coverslip) into a 15 ml falcon filled with 2.5 ml pancreatic islet medium. Centrifugation at 140 rcf for 3 min was followed by adding 1 ml pre-warmed trypsin (Trypsin-EDTA (0.05%), ref: 25300062, Gibco) to the pellet of cells. Next, the falcon was gently shaken in a water bath at 37 °C for 2 min. After adding 9 ml of islet medium and another centrifugation step (140 rcf, 3 min), the pellet was diluted in 500 µl – 2 ml medium, depending on the calculated cell number. Finally, the cells were harshly mixed and seeded in 50 – 100 µl droplets on coverslips. After 3 - 4 h incubation at 37 °C and 5% CO_2_, additional medium (2 ml for 6-well plate) was added to the attached cells.

### Homogenization of pancreatic islets

Homogenization of pancreatic islets was necessary for enzymatic assays and lipidomics sample preparation. It was achieved by initial dispersion of isolated pancreatic islets with trypsin and subsequent sonication (sonic dismembrator model 705, ref: 86853K-09-15, Thermo Fisher). After the final centrifugation step in the pancreatic islet dispersion, the pellet was reconstituted in Dulbecco’s phosphate-buffered saline (DPBS, ref: D8662, Merck/Sigma Aldrich) and the solution was sonicated at 50% for 2 min (20 s pulses and 20 s rest time) on ice.

### Glucose uptake measurement

Glucose uptake into pancreatic islets was assessed using the Glucose Uptake-Glo™ assay kit (ref: J1342, Promega). Groups of 5, 10 and 20 islets were washed in glucose-free Flex-medium (SILAC RPMI 1640 Flex Media, ref: A2494201, Gibco) and imaged for size normalization using a stereo microscope (Stereo microscope 305, ref: 3943002384, Zeiss) with a camera adapter (Axiocam 105 color). Afterwards, pancreatic islets were incubated for 1 h (37 °C and 5% CO_2_) in glucose-free Flex-medium with 20 mM of 2-Deoxyglucose (2DG, provided with the kit) and the standard protocol provided by the company was subsequently followed.

### Western Blot

Groups of 300 islets were collected in ice-cold PBS and centrifuged at 2000 rcf for 5 min (cooled down to 4 °C). The resulting pellet was snap-frozen in liquid nitrogen and stored at - 20 °C until further use. Cell lysis was performed for 30 min on ice using a cell lysis buffer (10 mM Tris-HCl, 10 mM NaCl, 0.1 mM EDTA, 0.5% Triton-X-100, 0.2% NaN_3,_ 200 µM PMSF, 1:100 protease/phosphatase inhibitor cocktail). Subsequently, the samples were centrifuged for 30 min at 14,000 rcf and 4 °C. The supernatant was transferred into a new reaction tube and protein concentration was assessed. The proteins were separated according to their molecular mass using sodium dodecyl sulfate-polyacrylamide gel electrophoresis (SDS-PAGE). Therefore, the samples were mixed with 2x Laemmli buffer (ref: S3401, Merck) in a 1:1 ratio and denatured for 5 min at 95 °C. Then, the denatured protein samples were added to a 10 or 12.5% SDS-polyacrylamide gel and separated at 100 V and 30 - 40 mA. The protein marker peqGOLD IV (ref: 27-2110, VWR) was used to quantify protein size. Before the separated proteins could be visualized with antibodies, they first had to be transferred to a methanol-activated polyvinylidene difluoride (PVDF) membrane using the semi-dry approach. Here, the SDS gel and the PVDF membrane were packed between Whatman papers in a semi-dry chamber. After applying 25 V and 1.3 A for 7 min, the membrane was taken out of the chamber and unspecific binding sites were blocked using Tris-buffered saline with 0.1% Tween^®^ 20 detergent (TBST) and 5% BSA for 1 h at RT. Primary antibodies were diluted 1:500 in TBST with 1% BSA and incubated with the membrane overnight at 4 °C. On the next day, membranes were washed 3 times for 10 min with TBST before incubation with HRP-conjugated secondary antibody for 1 h at RT. After another washing step with TBST (3 times for 10 min), the enhanced chemiluminescence (ECL) solution (ref: 1705060, Bio-Rad was added to visualise the proteins on the membrane. Western blot images were acquired with a gel documentation system (Gel Doc XR+, Bio-Rad, Germany), analyzed using the Image Lab software and normalized to the housekeeping protein (β-actin, Suppl. Table 1).

Presentation and statistical analysis of WB data were performed in a paired manner. One *Taz*- KD sample was always compared to another identically prepared WT sample. Therefore, the WT samples are all displayed at 100% and do not show an error among them. However, the variation among the samples is presented in separated figures in the supplements. The reason being that depending on the functional experiments some samples were frozen after 1 day and others after 2 or 3 days in culture.

### Glucokinase assay

The fluorometric glucokinase activity assay kit (ab273303) was used to study glucokinase activity. If not stated otherwise, the protocol provided by the manufacturer was followed. Groups of 150 freshly isolated pancreatic islets were dispersed and homogenised. Background intensity was measured for each sample. Fluorescent intensity (λ_ex_ = 540/20 nm, BS = 560 nm, λ_em_ = 590/20 nm) was measured using a Clariostar plate reader (BMG) and the obtained results were normalized by BCA protein assay.

### Glucose-6-phosphate dehydrogenase (G6PDH) enzymatic assay

Activity of glucose-6-phosphat dehydrogenase was assessed using a fluorometric kit from abcam (ab176722). Groups of 50 pancreatic islets were dispersed and homogenized. The protocol was performed according to the guidelines of the manufacturer. Fluorescent intensity (λ_ex_ = 535/20 nm, BS = 561 nm, λ_em_ = 587/20 nm) was measured using a Clariostar (BMG) plate reader and normalized to protein levels.

### Measurement of the mitochondrial oxygen consumption rate and extracellular acidification rate

Oxygen consumption (OCR) and extracellular acidification rate (ECAR) of whole pancreatic islets were assessed using a Seahorse XFe96 Analyzer (Agilent). One day before the experiments, sensor cartridges were prepared by adding calibrant XF (100840-000, Agilent) and incubating overnight at 37 °C (no additional CO_2_). On the day of the experiment, a spheroid XFe96 microplate (102978-100, Agilent) was coated with poly-L-lysine, and each well was filled with 175 µl of pre-warmed Seahorse XF RPMI medium (103576-100, Agilent) supplemented with 0.1% FBS, 2.8 mM glucose (103577-100, Agilent), and 2 mM glutamine (10359-100, Agilent). Groups of 15 islets were seeded with 5 µl volume into the corresponding detent at the bottom of the wells (29) and the plate was equilibrated for 1 h in a non-CO_2_ incubator. Afterwards, the protocol of the Seahorse XF Cell Mito Stress Test Kit (103015-100, Agilent) combined with an initial glucose stimulation (2.8, 10, or 20 mM) was conducted. The concentrations for the inhibitors were optimised in initial islet experiments and set to 4.5 µM of oligomycin, 1 µM of FCCP, 5 µM of antimycin A and rotenone. The assay was performed with at least six replicates per condition. Samples that did not match the following criteria were excluded: Absolute oxygen concentration between 130 - 160 mmHg and stable throughout the assay. The spheroid plate was inspected after the measurement to see if the islets were still inside the small detent of each well. Analysis was done with Wave software (version 2.6.3) and Prism (version 9.4).

Nutrient dependencies and capacities were measured using the Seahorse XF Mito Fuel Flex Test Kit (103260-100, Agilent). In detail, three inhibitors (4 µM of etomoxir, 2 µM of UK5099 and 3 µM of BPTES) were used to test each genotype’s nutrient dependencies and capacities. First, only one pathway was blocked by the injection of one inhibitor and subsequently, the other two pathways were blocked together with the next injection. Different combinations of inhibitors could determine the nutrient dependencies and capacities. The Seahorse assay medium for this nutrient dependency test was based on the Seahorse XF RPMI medium supplemented with 10 mM glucose, 2 mM glutamine, 1 mM pyruvate (103578-100, Agilent), and 0.1% FBS.

### ATP assay

ATP content was determined by the CellTiter-Glo^®^ luminescent cell viability assay (ref: G7570, Promega). Initially, groups of 5, 10 and 20 pancreatic islets were formed under standard culture conditions and imaged using a stereo microscope with a camera adapter (for normalization by pancreatic islet size). Afterwards, the protocol provided by the manufacturer was followed and the luminescence (λ_em_ = 545/50 nm) was recorded with a Clariostar (BMG) plate reader. The luminescence signal was converted to ATP levels via a standard curve acquired from different concentrations (10 nM – 10 µM) of adenosine 5′-triphosphate disodium salt hydrate. In long-term experiments the pancreatic islets were pretreated with various substances: 1 µM Auranofin, 10 µM GLX351322 (GLX), exogenous H_2_O_2_ (10, 20 and 100 µM).

### SypHer pH measurements

The ratiometric SypHer pH sensor was used to test cytosolic pH levels in dispersed pancreatic islets. Adenoviral transduction was performed by adding 0.5 µl of adenovirus on top of a coverslip with 10 dispersed pancreatic islets in 2 ml RPMI medium (10% FBS, 1% P/S). 2 days after transduction, cells were measured (λ_ex1_ = 405/20 nm, λ_ex2_ = 470/40 nm, BS = 505 nm, λ_em_ = 550/100 nm) using an inverted epifluorescence microscope Axio Observer 7 (Zeiss) and a 20x objective (Zeiss). Different glucose conditions (2 mM and 20 mM) were applied, followed by the addition of 5 µM oligomycin, 2 µM FCCP and 30 mM NH_4_Cl.

### Mitochondrial membrane potential measurement

The dye tetramethylrhodamine methyl ester (TMRM) was used in quenching mode to measure the mitochondrial membrane potential of whole pancreatic islets. Quenching mode of TMRM refers to the massive accumulation of TMRM molecules inside the mitochondria. Accumulation can lead to changes in the observed spectral properties which can be detected with a plate reader by monitoring the fluorescence (λ_ex_ = 535/20 nm, BS = 557 nm, λ_em_ = 585/30 nm). The isolated pancreatic islets were incubated at 37 °C for 45 min in low glucose KHB (2 mM glucose) together with 200 nM of TMRM. After loading, the islets were washed and transferred in groups of 25 per well into a 96-well plate filled with low glucose KHB. The baseline was measured for 15 min and then, 20 mM of glucose (or 100 µM of tolbutamide) was applied. As a positive control, 25 µM of CCCP was added at the end of each experiment.

### Cytosolic calcium measurements

Intracellular cytosolic calcium levels of dispersed and whole pancreatic islets were measured using the ratiometric fluorescent dye Fura-2 AM (λ_ex1_ = 340/30 nm, λ_ex2_ = 387/15 nm, BS = 409 nm λ_em_ = 510/90 nm). Groups of 10 similar size WT and *Taz*-KD islets were loaded for 2 h with 5 µM Fura-2 AM under standard culture conditions. After washing and starvation with 2 mM glucose in KHB for 15 min, WT and *Taz*-KD islets were placed in the measurement chamber of the Axio Observer 7. Baseline was recorded for 10 min and then, glucose concentration was raised to 20 mM glucose. KCl (30 mM) was used as a positive control at the end of the experiment. In another set of experiments, tolbutamide was used in titration protocol (1 – 100 µM) in the presence of 2 mM glucose.

### Mitochondrial calcium measurements

The Mito-Pericam calcium sensor was used to test mitochondrial calcium concentration in dispersed pancreatic islets (30, 31). Adenoviral transduction was performed by adding 0.5 µl of Mito-Pericam adenovirus on top of a coverslip with 25 dispersed pancreatic islets in 2 ml RPMI medium (10% FBS, 1% P/S). Two to three days after transduction, cells were measured (λ_ex_ = 405/20 nm, BS = 505 nm, λ_em_ = 550/100 nm) using an inverted epifluorescence microscope Axio Observer 7 (Zeiss) and a 63x oil objective (Zeiss). The baseline was recorded for 5 min in KHB with 2 mM glucose, before a 20 mM glucose solution (final concentration) was applied. After 10 - 20 min measurement in high glucose conditions, 30 mM of KCl was applied to have a depolarizing control.

### Calcium measurement in ER-lumen

The ER-targeted biosensor D4ER was used to measure calcium levels in the ER lumen. A rat insulin promotor linked to the D4ER gene allowed specific transfection only in pancreatic β- cells (32). Groups of 25 dispersed islets were seeded on to coverslips in 2 ml RPMI medium (10% FBS, 1% P/S) and transduced by 0.5 µl of adenovirus encoding D4ER. 2 - 3 days after transduction, cells were measured (λ_ex_= 445/25 nm, BS = 505 nm, λ_em1_ = 475 nm, λ_em2_ = 540/20 nm) using an inverted epifluorescence microscope Axio Observer 7 (Zeiss) and a 63x oil objective (Zeiss). The baseline was recorded for 5 min in KHB with 2 mM glucose, before a 20 mM glucose solution (final concentration) was applied. At the end of the experiment, 3 µM thapsigargin (final concentration) was added to empty the ER calcium storage.

### Redox histology

Mito-roGFP2-Orp1 crossbreed with shTaz mice (MiOxTaz mice) were sacrificed via 2:1 Ketamin/Rompun (1 g/kg Ketavet^®^/ 100 mg/kg Rompun^®^) injection. Redox histology was performed as previously described with some modifications (23, 33). First, cardiac perfusion with 25 ml of 50 mM N-Ethylmaleimide (NEM) diluted in PBS was used to block free thiols and prevent artificial sensor oxidation. Afterwards the pancreas was inflated with 5 ml of the same solution. The whole pancreas was removed and fixation was performed overnight at RT with 4% PFA. To improve cryoslice quality, pancreas samples were additionally treated with 30% sucrose diluted in PBS for 3 h before embedding in Tissue Tek. All samples were cut at 5 µm using a cryostat at -15 °C. Pancreas slides were dried for 30 min before being washed with PBS and treated with 3% goat serum for 1 h in a wet chamber. Subsequently, 50 µl of primary anti-insulin antibody (Suppl. Table 1, 1:200 diluted) was added and incubated overnight in a wet chamber at 4 °C. On the following day, slides were again washed with PBS and 50 µl of Alexa 594 secondary antibody (Suppl. Table 1, 1:400 diluted) was added for 1 h 15 min at RT. After a final washing step, slides were dried and mounted with Dako mounting media. Imaging was performed with the previously described Axio Observer 7 system using a 20x air objective. The roGFP2-Orp1 fluorescence was measured at 500 to 550 nm with either 405/20 nm or 470/40 nm excitation. Pancreatic islets inside the pancreas tissue were localised using a mCherry filter set. Analysis was performed on ImageJ using a self-written code to automate the procedure and minimize risk of bias in setting thresholds.

After calculating the ratio for each image, both genotypes (WT and *Taz*-KD) were compared by calculating the percentage change:

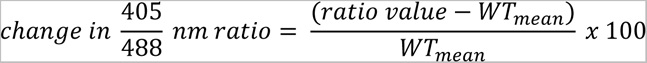

ratio value: either WT or *Taz*-KD ratio value; WT_mean_: mean value for all the WT ratio values.

### H_2_O_2_ measurements

After culture, islets were collected, washed in KHB containing 10 mM glucose and 0.1% BSA for 10 min, and transferred to a U-shaped 96-well plate (TPP, ref: 92097) at 20 - 25 islets per well in a total volume of 160 μl. roGFP2 fluorescence (λ_ex1_ = 400/10 nm λ_ex2_ = 482/16 nm, λ_em_ = 530/40 nm) was measured at 37 °C, 18% O_2_ and 5% CO_2_ for 19 h, using a microplate reader Clariostar (BMG Labtech).

### NADH/NADPH (NAD(P)H) autofluorescence

The NAD(P)H levels of WT and *Taz*-KD pancreatic islets were measured in Clariostar plate reader experiments (λ_ex_ = 340/10 nm, BS = 410 nm, λ_em_ = 450/10 nm) in parallel to other parameters (mitochondrial membrane potential and H_2_O_2_). In the H_2_O_2_ experiment, baseline NAD(P)H autofluorescence at 10 mM glucose was monitored over time at 37 °C, 18% O_2_ and 5% CO_2_. In the TMRM experiments, NAD(P)H autofluorescence was monitored in 2 mM and 20 mM glucose conditions and later normalized to CCCP levels.

### Confocal and STED microscopy

Directly after islet isolation, groups of 50 pancreatic islets were dispersed and seeded on a coverslip (170 µm thickness, # 1.5, Marienfeld, Gemany). On the next day, the dispersed islet cells were washed with 10 mM glucose KHB and stained with 30 nM MitoTracker^TM^ Deep Red in RPMI 1640 (only 0.1% FCS) for 10 min at RT. After three washing steps, the imaging was carried out in 10 mM glucose KHB at RT. The samples were imaged on an inverted STED microscope (Expert Line, Abberior Instruments, Göttingen, Germany) using the according control software Imspector V16.3. A 100x silicon immersion objective with a numerical aperture of 1.4 (UPLSAPO100XS, Olympus, Hamburg, Germany) and a pinhole size of 90.0 µm (1.08 airy units) was used. In order to have an overview of the pancreatic islet cells with the labeled mitochondria, single confocal 80 x 80 µm scanning with 200 nm (in XY axis) was performed. at excitation of 640 nm. This was followed by another confocal scanning of a randomly selected cell, this time with finer XY pixel size (40 nm) and a Z-stack to generate a 3D image (voxel size: 40 x 40 x 300 nm^3^). The detection filter and total pixel dwell time were set to 650 – 720 nm and 5 µs, respectively. Finally, parts of the cellular volume were recorded in STED mode. The STED experiments were performed using “rescue settings” which reduce the photobleaching of the Mitotracker. The 775 nm depletion laser pulsed at 40% of the maximal power of 1250 mW (corresponding to 75 – 85 mW in the focus, repetition rate of 40 MHz) with no gating and total dwell time of 17.5 µs. The STED images were recorded with a voxel size of 20 x 20 x 300 nm³. In addition to confocal and STED image stacks, the residual excitation by the STED beam (so called “re-excitation”) was recorded and later subtracted from the STED recordings after deconvolution.

Data were plane-wise linearly deconvoluted using a Wiener filter with theoretical point spread functions and manually adjusted regularisation parameter. For deconvolution and subtraction of the re-excitation custom-written routines in MATLAB were used. Next, all images were preprocessed in ImageJ for feature extraction using smoothing and shot-noise reduction (bilateral filter), background subtraction (rolling-ball algorithm) and sharpening (unsharp mask). Subsequently, threshold-based image segmentation and 3D-rendering was performed in Imaris (version 9.6). The resulting disconnected surfaces were grouped based on their surface area into three classes: Class A = 0.3 - 3 µm^2^, Class B = 3 - 10 µm^2^, and Class C > 10 µm^2^. The morphological parameters, including surface area, volume, number of single mitochondria, sphericity, and bounding boxes were automatically calculated by Imaris software based on each disconnected object. The sphericity describes how spherical an object is and can be calculated via 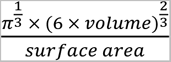. The bounding box of a single mitochondrion describes the minimal rectangular box which fully encloses the object. Furthermore, a nearest neighbor analysis was performed with a custom-written MATLAB protocol. The program gives the number of neighboring objects for each object in a radius of 2.5 µm.

### Mitotracker for mitochondrial volume

MitoTracker^TM^ Deep Red (M22426, Thermo Fisher) was used to visualize and measure mitochondrial volume. To quantify the mitochondrial volume, MitoTracker signal (λ_ex1_ = 620/60 nm, BS = 665 nm, λ_ex1_ = 700/30 nm) was normalized by the Hoechst 33342 (62249, Thermo Fisher) signal. For staining, groups of 50 pancreatic islets were dispersed. The dispersed pancreatic islet cells were washed with KHB and incubated with 50 nM Mitotracker for 30 min at 37 °C in RPMI supplemented with only 0.1% FCS. After two washing steps in KHB, the cells were incubated for 7 min with 5 µg/ml Hoechst dye in KHB (0.1% BSA & 10 mM glucose) at RT protected from light. Before imaging in KHB, the cells were washed three times in KHB (0.1% BSA & 10 mM glucose).

### Citrate synthase assay

The colorimetric Citrate Synthase Assay Kit (ab239712) was used to determine the mitochondrial mass. A Clariostar plate reader was used to measure the absorbance (λ_ex_ = 412 nm). The citrate synthase activity was normalized by protein content.

### RNA Sequencing

RNA samples were diluted in RNase-free water and 5 µl of RNA sample were sent for sequencing, which was performed by Novogene (Novogene GmbH, Cambridge, UK). The quantity and quality of the RNA samples were assessed using the following methods. Preliminary quality control was performed on 1% agarose gel electrophoresis to test RNA degradation and potential contamination. Sample purity and preliminary quantitation were measured using Bioanalyzer 2100 (Agilent Technologies, USA) and it was also used to check the RNA integrity and final quantitation.

For library preparation, we used the Novogene NGS RNA Library Prep Set (PT042). The mRNA present in the total RNA sample was isolated with magnetic beads of oligos d(T)25. This method is known as polyA-tailed mRNA enrichment. Subsequently, mRNA was randomly fragmented and cDNA synthesis proceeded using random hexamers and the reverse transcriptase enzyme. Once the synthesis of the first chain is finished, the second chain is synthesized with the addition of an Illumina buffer (non-directional library preparation). With this and together with the presence of dNTPs, RNase H and polymerase I from *E. Coli*, the second chain will be obtained by Nick translation. The resulting products go through purification, end-repair, A-tailing and adapter ligation. Fragments of the appropriate size are enriched by PCR, where indexed P5 and P7 primers are introduced, and final products are purified.

The library was checked with Qubit 2.0 and real-time PCR for quantification and bioanalyzer Agilent 2100 for size distribution detection. Quantified libraries were pooled and sequenced on the Illumina Novaseq X platform, according to effective library concentration and data amount using the paired-end 150 strategy (PE150).

### Statistical Analysis

GraphPad Prism Software version 9.4 was used for statistical analysis. The presented values are shown in mean ± SEM. Details on the statistical analysis are in the figure legends.

## Results

### *Taz*-KD alters CL levels and lipid species profile

To decrease Taz1 levels we used male and female mice expressing a short hairpin RNA (shRNA) to attenuate *Taz* expression (*Taz*-KD mice). *Taz*-KD mice and WT littermates were fed with doxycycline (doxy) to either induce *Taz* knockdown or serve as a control (Fig. 1A, Fig. Suppl. 1A). *Taz*-KD mice were smaller and had decreased body weight at 20 weeks (Fig. 1B, C) and 50 weeks, but not at 10 weeks of age (wo) (Fig. Suppl. 1B) compared to WT littermates. Reduced *Taz* expression was confirmed in hearts and isolated pancreatic islets from 20 wo *Taz*-KD mice. Decrease in *Taz* expression was more pronounced in the heart (∼90% decrease) than in pancreatic islets (∼60% decrease) (Fig. 1D, E and Fig. Suppl. 1C, D).

**Figure 1:**
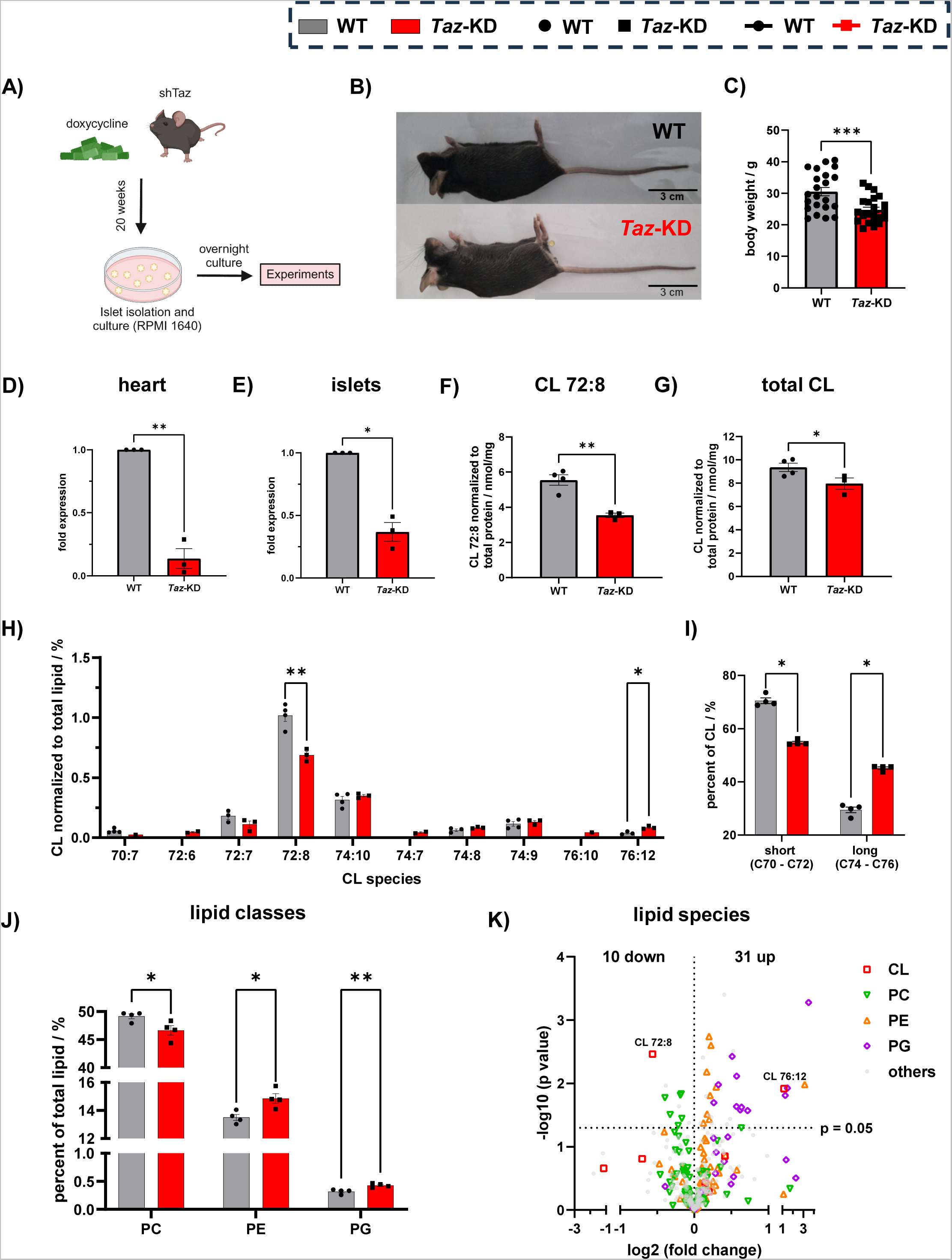
CL reduction and phospholipid alterations in *Taz*-KD pancreatic islets. (**A**) Schematic illustration of shTaz mouse model with 20 weeks of doxy feeding. Pancreatic islets are isolated, cultured in RPMI 1640 with 10% FBS and 1% P/S and experiments are performed after overnight culture. Illustration is created with Biorender.com. (**B**) Representative image of a 20 wo *Taz*-KD mouse compared to a WT littermate. (**C**) Body weight of *Taz*-KD and WT at 20 wo, N = 22. *Taz* gene expression in heart (**D**) and pancreatic islet (**E**) tissue. N = 3. CL levels of the main species CL 72:8 (**F**) and total CL amount (**G**) of *Taz*-KD and WT pancreatic islets at 20 wo normalized to protein concentration. (**H**) CL species profile of *Taz*-KD and WT pancreatic islets at 20 wo normalized to total lipid amount, N (WT) = 4, N (*Taz*-KD) = 3, some replicates are below the limit of detection. (**I**) Quantification of the acyl chain length of all CL species (short: C70 - C72 and long: C74 - 76), N = 4. (**J**) Lipid concentration of the phospholipids PC, PE and PG of *Taz*-KD and WT pancreatic islets at 20 wo normalized to total lipid amount, N = 4. The whole lipid class profile is presented in the supplements (Fig. Suppl. 1F). (**K**) Volcano plot of all detected lipid species of *Taz*-KD and WT pancreatic islets at 20 wo. Statistical analysis showed that 10 lipid species are significantly (p < 0.05) downregulated and 31 are upregulated in *Taz*-KD. The lipid classes CL, PC, PE, and PG are highlighted. All significantly altered lipid species are listed in the supplements (Fig. Suppl. 1F). Data represent mean ± SEM (indicated by error bars); N numbers indicate number of animals; statistical significance was determined by unpaired Student *t* test: *p < 0.05, **p < 0.01, ***p < 0.001. Abbreviations: weeks of age (wo), *Tafazzin*-Knockdown (*Taz*-KD), Wildtype (WT), *Tafazzin* (*Taz*), Cardiolipin (CL), Phosphatidylcholine (PC), Phosphatidylethanolamine (PE), Phosphatidylglycerol (PG).

We first assessed the impact of decreased *Taz* expression on the lipid profile of pancreatic islets. To this end, we performed a lipidomics analysis on pancreatic islets isolated from 20 wo *Taz*-KD and WT mice. As expected, the main CL species tetralinoleoyl cardiolipin (CL 72:8) and total CL levels were significantly decreased when normalized to total protein content (Fig. 1F, G). In addition, the relative abundance of different CL species (normalized to total lipid amount) was altered in pancreatic islets from *Taz*-KD mice compared to WT littermates (Fig. 1H). We found a decrease in CL species with shorter acyl-chain lengths and an increase in CL species with longer acyl-chain lengths (Fig. 1I). Furthermore, the full lipid class profile of *Taz*- KD mice revealed a reduction of phosphatidylcholines (PC) and an increase in phosphatidylethanolamines (PE) and phosphatidylglycerols (PG), which are involved in CL biosynthesis and remodeling (34) (Fig. 1J and Fig. Suppl. 1E). In total, we observed a significant decrease of 10 and increase of 31 lipid species in *Taz*-KD pancreatic islets as shown in the volcano plot, highlighting the most relevant lipid classes (Fig. 1K and Fig. Suppl. 1F). In summary, knockdown of the transacylase *Taz* affects the lipidome of pancreatic islets, particularly altering the CL profile, reducing CL content, and changing PC, PE, and PG content.

### *Taz*-KD increases glucose uptake, respiratory capacity in low glucose with no changes in ATP production

To investigate whether changes in CL and lipid profile would impact the function of *Taz*-KD pancreatic islets, we systematically investigated several steps of glucose metabolism, which culminate in insulin secretion. Briefly, high capacity, low affinity, plasma membrane glucose transporters (mainly GLUT2 in mouse pancreatic β-cells) allow the rate of glucose influx to increase proportionally to increases in blood glucose level. Glucose is subsequently predominantly metabolized via the glycolytic pathway and TCA cycle, ultimately leading to respiratory chain ATP production. The flux through these pathways and thus the rate of ATP production responds directly and proportionally to changes in blood glucose levels (35)( (Fig. 2A). Islets isolated from 20 wo *Taz*-KD mice showed increased glucose uptake compared to WT pancreatic islets (Fig. 2B), which was not due to an increase in GLUT2 protein level (Fig. 2C, and Fig. Suppl. 2A). Interestingly, we also observed that basal (Fig. Suppl. 2B) and glucose-stimulated NAD(P)H levels (measured by autofluorescence) were increased in *Taz*- KD islets compared to WT (normalized to NAD(P)H levels measured in islets treated with the membrane uncoupler, CCCP) (Fig. 2D). Increased NAD(P)H levels were independent of glucokinase (GCK) and glucose-6-phosphate-dehydrogenase (G6PDH) activity, which was not different between *Taz*-KD and WT islets (Fig. 2E, F).

**Figure 2:**
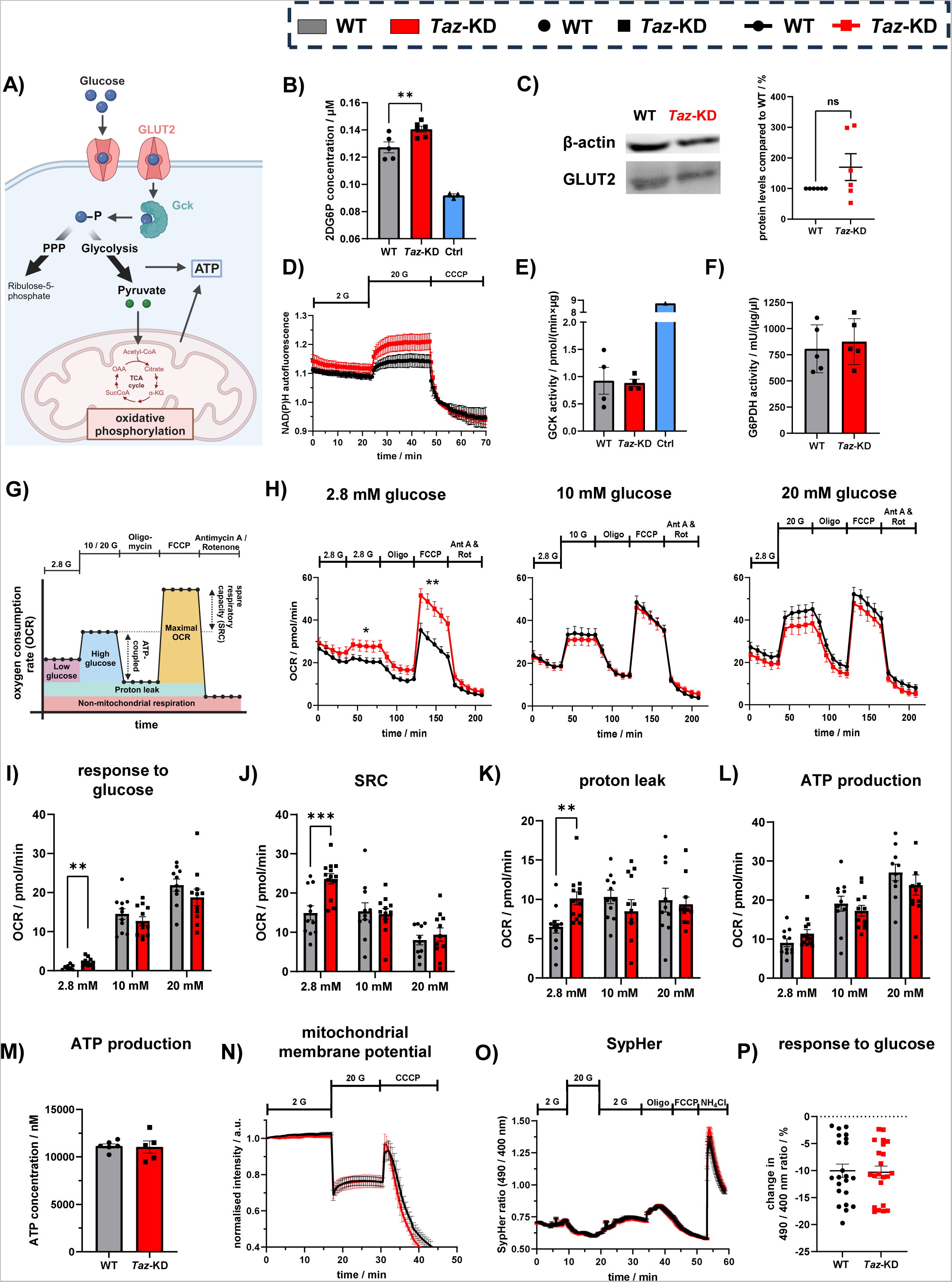
Increased glucose uptake and amplified metabolic parameters in *Taz*-KD pancreatic islets with no effects on ATP levels. (**A**) Schematic figure of glucose metabolism in pancreatic islet cells. Created with Biorender.com. (**B**) Glucose uptake of 20 wo WT and *Taz*-KD pancreatic islets, indicated by the levels of 2DG6P. Pancreatic islets without 2DG loading were used as negative controls (blue). N (WT) = 5, N (*Taz*-KD) = 6, N (Ctrl) = 3. (**C**) Representative western blot (left) and quantification (right) of GLUT2 normalized to β-actin in pancreatic islets of 20 wo *Taz*-KD mice, N = 6. (**D**) NAD(P)H autofluorescence measurement of 20 wo WT and *Taz*-KD pancreatic islets in 2 mM and 20 mM glucose, normalized to CCCP, N (WT) = 4, N (*Taz*-KD) = 6. Quantification of GCK (**E**) and G6PDH (**F**) enzyme activity. A positive control for GCK activity was provided by the kit (blue). N (GCK) = 4, N (G6PDH) = 5. (**F**) (**G**) Schematic protocol of OCR and quantifiable parameters during mitochondrial stress test using Seahorse created with Biorender.com. (**H**) OCR kinetics of 20 wo WT and *Taz*-KD pancreatic islets in response to 2.8 mM (left), 10 mM (middle) and 20 mM (right) glucose stimulation followed by the addition of inhibitors of the respiratory chain complexes (Oligo, Ant A and Rot) and uncoupler (FCCP). n (WT) = 11, n (*Taz*-KD) = 13. Quantification of response to glucose (**I**), SRC (**J**), proton leak (**K**), ATP production (**L**) separated by glucose concentrations (2.8, 10 and 20 mM), n (WT) = 11, n (*Taz*-KD) = 13, n number of experiments include N (WT) = 5 and N (*Taz*-KD) = 4. (**M**) ATP concentration of 20 wo WT and *Taz*-KD pancreatic islets using the CellTiter-Glo^®^ assay, N (WT) = 6, N (*Taz*-KD) = 5. (**N**) Kinetic measurement of mitochondrial membrane potential of 20 wo WT and *Taz*-KD pancreatic islets using TMRM in quenching mode (200 nM) in presence of 2 and 20 mM glucose. To uncouple the mitochondria 25 µM of CCCP was added as a control at the end of the experiment. n = 16. (**O**) Cytosolic pH measurement of 20 wo WT and *Taz*-KD pancreatic islets transduced with SypHer adenovirus, following the change in SypHer ratio in the presence of 2 or 20 mM glucose, oligo, and FCCP. NH_4_Cl was added as a positive control in the end of the experiment. n (WT) = 7, n (*Taz*-KD) = 6, n number of experiments include N = 3 (number of animals). (**P**) Quantification of change in SypHer ratio in response to glucose (2 to 20 mM), n = 22, n number of experiments include N = 4 (number of animals). Data represent mean ± SEM (indicated by error bars); N numbers indicate number of animals; statistical significance was determined by unpaired Student *t* test: *p < 0.05, **p < 0.01, ***p < 0.001. Abbreviations: weeks of age (wo), *Tafazzin*-Knockdown (*Taz*-KD), Wildtype (WT), 2- deoxy-D-glucose (2DG), 2-deoxy-D-glucose-6-phosphate (2DG6P), glucokinase (GCK), glucose-6-phosphate dehydrogenase (G6PDH), oxygen consumption rate (OCR), oligomycin (Oligo), antimycin A (Ant A), rotenone (Rot), spare respiratory capacity (SRC).

Next, we used a Seahorse XFe96 Analyzer to study changes in oxygen consumption rate (OCR) and extracellular acidification rate (ECAR) upon stimulation with glucose (2.8, 10 and 20 mM) in isolated WT and *Taz*-KD pancreatic islets from 20 wo mice. Using an adapted protocol of the Seahorse Cell Mitochondrial Stress Test (Fig. 2G), we quantified various metabolic parameters. Under low glucose conditions (2.8 mM glucose), we observed an increased OCR in *Taz*-KD pancreatic islets (Fig. 2H), an increased response to glucose, increased spare respiratory capacity (SRC), increased proton leak and increased maximum respiration (Fig. 2H–K, Fig. Suppl. 2C). No differences in these parameters were seen with 10 and 20 mM glucose (Fig. 2 H–K). Basal (unstimulated) and non-mitochondrial respiration were unchanged between *Taz*-KD and WT islets (Fig. Suppl. 2D). Unexpectedly, ATP-coupled OCR was not different in *Taz*-KD pancreatic islets (Fig. 2L). A luminescence-based ATP assay confirmed that ATP levels are not different between *Taz*-KD and WT islets after stimulation with 10 mM glucose (Fig. 2M). To investigate if pancreatic islet mitochondrial metabolism is dependent on other metabolic substrates, as observed in other tissues from *Taz*-KD models (for example heart and patient iPSC cell-derived cardiomyocytes) (9, 14), we performed the Seahorse XF Mito Fuel Flex test, where inhibitors for the mitochondrial pyruvate carrier (UK5099), lipid β-oxidation (etomoxir), and glutamine metabolism pathway (BPTES) are employed (Fig. Suppl. 2I). Apart from a slight increase in OCR of *Taz*-KD islets after BPTES addition (Fig. Suppl. 2J), we did not observe any differences when comparing WT and *Taz*-KD pancreatic islets neither in terms of nutrient preferences (Fig. Suppl. 2J), nor in nutrient capacities by inhibiting all the other metabolic pathways (Fig. Suppl. 2K). In line with the OCR results, we did not observe any difference in the mitochondrial membrane potential of *Taz*-KD and WT pancreatic islets using the membrane potential fluorophore TMRM in quenching mode (Fig. 2N). Another parameter of cellular metabolism acquired during the Seahorse experiments is the ECAR. We did not observe alterations in basal and glucose-stimulated ECAR levels in *Taz*-KD pancreatic islets (Fig. Suppl. 2E). Furthermore, we observed no difference in ECAR dynamics in either the mitochondrial stress test assay (Fig. Suppl. 2F-H), or the Mito Fuel Flex test (Fig. Suppl. 2L, M) between *Taz*-KD and WT pancreatic islets. Of note, in most cell types, the addition of oligomycin leads to an increase in anaerobic glycolysis and ECAR. However, we observed diminished ECAR for pancreatic islets treated with oligomycin, which is consistent with a previous report in Min6 pseudo islets (36). Decreased ECAR after oligomycin treatment might be connected to low LDH expression levels and low lactate production in pancreatic β- cells (37). We could confirm the finding of a similar acidification rate in *Taz*-KD and WT pancreatic islets by employing the genetically encoded cytosolic pH sensor SypHer, which reported similar pH responses to glucose and respiratory chain inhibitors (Fig. 2O, P). In summary, *Taz* knockdown increased pancreatic islet glucose uptake and several parameters of cellular metabolism in low glucose including SRC, proton leak, and maximal respiration, without leading to any change in ATP level.

### *Taz* knockdown leads to faster calcium influx

To gain insight into the downstream impacts of increased glucose uptake in *Taz*-KD islets, we investigated several parameters. An increase in the ATP/ADP ratio in pancreatic β-cells in response to increased blood glucose level leads to the closure of plasma membrane K_ATP_ channels, resulting in membrane depolarization and calcium influx, which ultimately triggers insulin secretion (Fig. 3A). Interestingly, although we observed similar cytosolic calcium levels in *Taz*-KD and WT pancreatic islets under both low and high glucose conditions (Fig. 3B, Fig. Suppl. 3A), *Taz*-KD pancreatic islets displayed a faster calcium influx upon glucose stimulation (Fig. 3B, C). To test if faster calcium response was based on an altered threshold for channel opening, we performed titration experiments with the K_ATP_ channel blocker tolbutamide. The calculated dose-response curve showed a tendency towards increased sensitivity of calcium influx rate to tolbutamide in *Taz*-KD pancreatic islets, although no statistically significant difference was found (Fig. Suppl. 3B, C). The calcium handling inside other organelles plays a crucial role in regulating the cytosolic calcium concentration upon stimulation with glucose (30, 32, 38). Therefore, we dispersed WT and *Taz*-KD pancreatic islets and used adenoviral vectors to express ER- and mitochondrial matrix-targeted calcium sensors to investigate calcium handling in these cellular compartments in *Taz*-KD cells (Fig. 3D). The response of mitochondrial (Fig. 3E) and ER calcium (Fig. 3F) to glucose stimulation was very similar in *Taz*- KD and WT pancreatic islet cells. Finally, we investigated the cytosolic calcium response in dispersed pancreatic islet cells of WT and *Taz*-KD. Interestingly, the faster cytosolic calcium influx observed in whole *Taz*-KD pancreatic islets was absent in dispersed *Taz*-KD pancreatic islet cells compared to WT (Fig. 3G).

**Figure 3:**
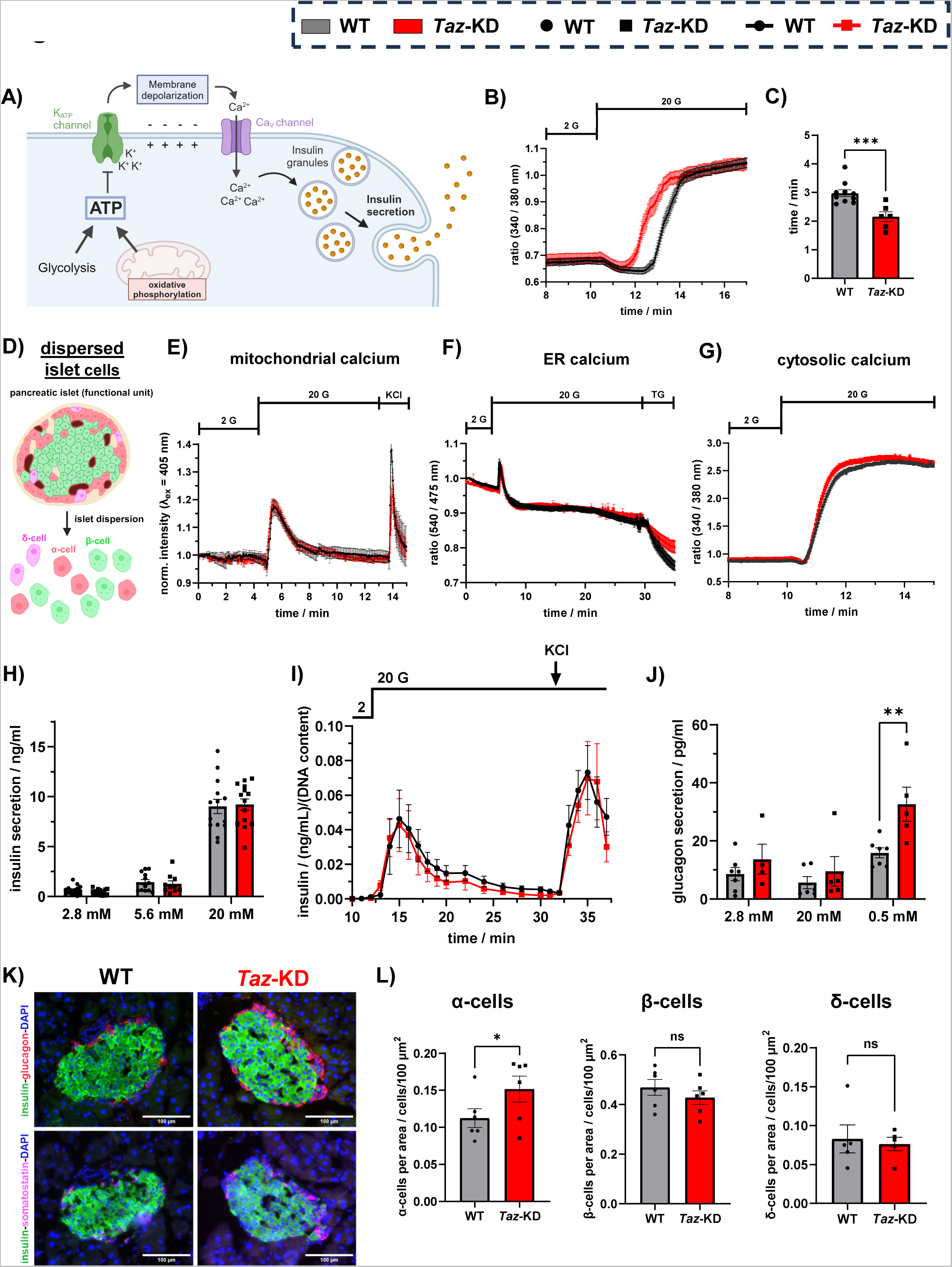
Faster cytosolic calcium influx and increased glucagon secretion in *Taz*-KD pancreatic islets. (**A**) Schematic illustration of ATP-dependent insulin secretion in pancreatic β-cells, created with Biorender.com. (**B**) Cytosolic calcium levels of 20 wo WT and *Taz*-KD pancreatic islets in 2 mM and 20 mM glucose concentration, monitored by Fura-2 AM ratio. (**C**) Quantification of the timing of the cytosolic calcium influx upon glucose stimulation (20 mM), N (WT) = 10, N (*Taz*-KD) = 6. (**D**) Schematic illustration of pancreatic islet dispersion from whole pancreatic islets that act as a functional unit to single islet cells with limited connection between the cells, created with Biorender.com. Mitochondrial (**E**), ER (**F**) and cytosolic (**G**) calcium levels of dispersed pancreatic islet cells from WT and *Taz*-KD 20 wo mice, measured with Mito-Pericam, D4ER and Fura-2 AM respectively. N (mitochondrial calcium, WT) = 4, N (mitochondrial calcium, *Taz*-KD) = 5, N (ER calcium, WT) = 4, N (ER calcium, *Taz*-KD) = 5, N (cytosolic calcium) = 3. (**H**) Quantification of static *ex vivo* GSIS of 20 wo WT and *Taz*-KD pancreatic islets at 2.8, 5.6 and 20 mM glucose concentrations, N (WT) = 16, N (*Taz*-KD) = 17. (**I**) Dynamic *ex vivo* GSIS of 20 wo WT and *Taz*-KD pancreatic islets normalized to DNA content, following the insulin levels at 2 mM glucose, 20 mM glucose and 30 mM KCl conditions, N (WT) = 4, N (*Taz*-KD) = 6. (**J**) Glucagon secretion of 20 wo WT and *Taz*-KD pancreatic islets in a stimulatory glucose-based secretion protocol, N (2.8 mM, WT) = 6, N (20 mM, WT) = 5, N (0.5 mM, WT) = 7, N (2.8 mM, *Taz*-KD) = 4, N (20 and 0.5 mM, *Taz*-KD) = 5. (**K**) Representative images of 20 wo WT (left) and *Taz*-KD (right) pancreas cryoslices IHC showing pancreatic islets stained against insulin (green) - glucagon (red) (top panel) or insulin (green) - somatostatin (magenta) (bottom panel) together with DAPI (blue). The glucagon- and somatostatin-insulin double stainings for each genotype represent the same pancreatic islet at a different cutting depth. Scale bar: 100 µm. (**L**) Quantitative ImageJ analysis of α- (left), β- (middle) and δ- (right) cell number of IHC on cryoslices and counting DAPI spots of WT and *Taz*-KD pancreatic islets at 20 wo normalized to pancreatic islet area, N = 6. Data represent mean ± SEM (indicated by error bars); N and n numbers indicate number of animals and experiments, respectively; statistical significance was determined by unpaired Student *t* test or two-way ANOVA for glucagon secretion: *p < 0.05, **p < 0.01, ***p < 0.001. Abbreviations: weeks of age (wo), *Tafazzin*-Knockdown (*Taz*-KD), Wildtype (WT), glucose-stimulated insulin secretion (GSIS), immunohistochemistry (IHC).

We then tested secretory function upon different glucose concentrations in isolated pancreatic islets from 20 wo WT and *Taz*-KD mice. Analysis of static glucose-stimulated insulin secretion (GSIS) revealed no difference in insulin secretion (Fig. 3H) and insulin content at a range of difference glucose concentrations (Fig. Suppl. 3D). Moreover, we performed a dynamic insulin secretion analysis to investigate the different phases of insulin secretion. We observed no difference in first- and second-phase insulin secretion dynamics between WT and *Taz*-KD pancreatic islets (Fig. 3I, Fig. Suppl. 3E) either following glucose stimulation or after islet depolarization.

### *Taz*-KD increases islet glucagon secretion and α-cell number

We finally asked whether *Taz* knockdown might lead to changes in glucagon secretion. To this end, we performed a stimulatory glucagon assay where we first challenged the pancreatic islets with high glucose (20 mM) and then exposed them to low glucose (0.5 mM) to induce glucagon secretion. Interestingly, we observed increased glucagon secretion in *Taz*-KD pancreatic islets (Fig. 3J), and a tendency towards increased islet glucagon content, although this difference was not statistically significant (Fig. Suppl. 3F). Due to the differences observed in glucagon secretion, we next asked whether pancreatic islet cell composition was changed in *Taz*-KD mice (Fig. 3K, Fig. Suppl. 3G). *Taz*-KD pancreatic islets showed an increased number of glucagon-positive α-cells (Fig. 3K, L left panel), but no difference in insulin-positive β-cells (Fig. 3K, L middle panel) and somatostatin-positive δ-cells compared to WT (Fig. 3K, L right panel). We also analyzed PDX1, an important pancreatic β-cell specific transcription factor. Immunofluorescence staining for PDX1 showed no difference between WT and *Taz*-KD pancreatic islets at 20 wo (Fig. Suppl. 3H). We next analyzed if markers for proliferation (Ki67) or apoptosis (cleaved caspase-3) were changed in α-cells or β-cells. The number of α- and β- cells found positive for Ki67 (Fig. Suppl. 3I) and cleaved caspase-3 (Fig. Suppl. 3J) were also similar in 20 wo WT and *Taz*-KD pancreatic islets.

In summary, *Taz*-KD pancreatic islets have faster calcium influx and increased α-cell number with higher glucagon secretion and preserved β-cell function (insulin secretion). Calcium dynamics in dispersed pancreatic islet cells were very similar in WT and *Taz*-KD, indicating that intercellular communication within the pancreatic islets plays an important role in the *Taz*- KD phenotype.

### *Taz*-KD does not lead to *in vivo* and *ex vivo* H_2_O_2_ production in pancreatic islets

BTHS patient tissue samples and cell line models have been reported to exhibit increased levels of reactive oxygen species (ROS), which were previously suggested to be causally related to the BTHS phenotype (11). On the other hand, we and others have previously shown that in the hearts of *Taz*-KD mice, ROS levels are unchanged due to an enhanced antioxidative defense system (8, 39). As pancreatic islets are generally considered to be sensitive to ‘redox stress’ due to the low levels of antioxidant enzymes (40) and as ROS have been proposed to have a role as a metabolic coupling factor for insulin secretion (41, 42), we sought to further investigate the production and effects of ROS on pancreatic islets in *Taz*-KD mice. For that, we crossbred *Taz*-KD and WT mice with mice ubiquitously expressing a mitochondrial matrix-targeted H_2_O_2_ sensor, roGFP2-Orp1 (mito-roGFP2-Orp1) (Fig. 4A). First, we confirmed that the resulting transgenic mice (shTaz x mito-roGFP2-Orp1) were expressing the mito-roGFP2- Orp1 sensor and characterized the sensor response to H_2_O_2_ (Fig. Suppl. 4A). Consistent with previous observations, the mito-roGFP2-Orp1 sensor responds sensitively to exogenous H_2_O_2_ applied at an initial concentration of 25 μM. Addition of 100 µM H_2_O_2_ leads to maximal roGFP2- Orp1 oxidation, while DTT leads to maximal reduction (Fig. Suppl. 4A). We then used a redox histology approach to analyze the *in vivo* mito-roGFP2-Orp1 oxidation in 20 and 50 wo *Taz*- KD pancreatic islets (Fig. 4B) (23, 43). Surprisingly, we observed a significant reduction in the mito-roGFP2-Orp1 sensor (Fig. 4C - D), demonstrating a decreased mitochondrial redox state in pancreatic islets of *Taz*-KD mice.

**Figure 4:**
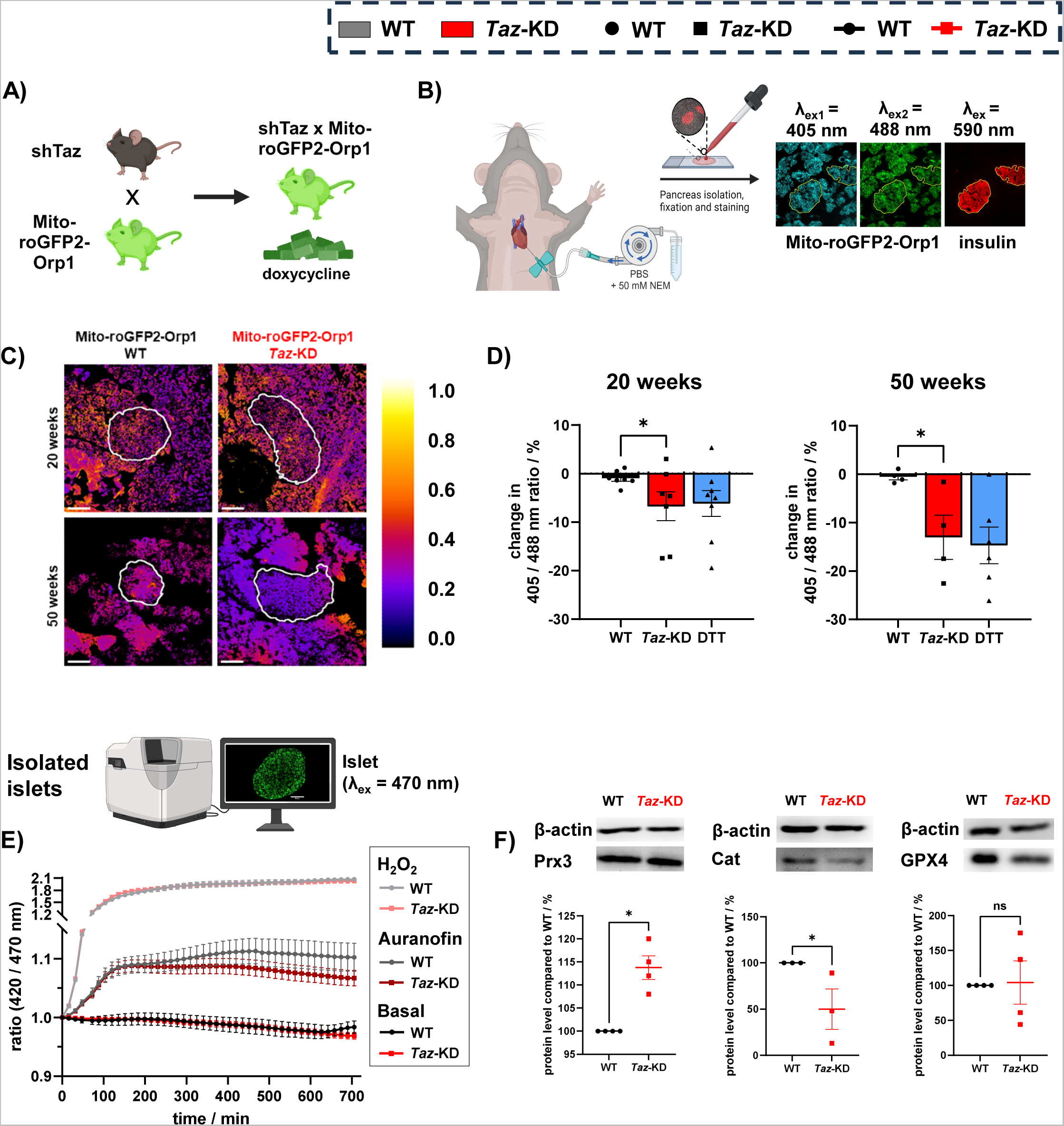
*Taz*-KD pancreatic islets exhibit reduced redox state and unchanged H_2_O_2_ dynamics. (**A**) Generation of a new mouse (shTaz x Mito-roGFP2-Orp1) model that expresses an shRNA against *Taz* and the mitochondrial H_2_O_2_ sensor roGFP2-Orp1. The mice are lifelong feed with doxy. Figure created with Biorender.com. (**B**) Schematic protocol of redox histology to measure *in vivo* redox state. Before pancreas isolation, shTaz x Mito-roGFP2-Orp1 mice are perfused with NEM via the cardiovascular system. Isolated pancreas is fixed in PFA und cryocuts are staining against insulin to localize the pancreatic islets. Imaged intensity ratio (excitation: 405/488 nm and emission: 500 – 530 nm) of the mito-roGFP2-Orp1 sensor in pancreatic islets reflects the *in vivo* redox status. (**C**) Representative ratiometric image (ImageJ Lookup table: “Fire”) of mito-roGFP2-Orp1/WT (left) and mito-roGFP2-Orp1/*Taz*-KD (right) pancreatic islets at 20 (top) or 50 wo (bottom). Scale bar: 100 µm. (**D**) Normalized percentage change in ratio of the redox state of the mito-roGFP2-Orp1 sensor in pancreatic islets of 20 (left) and 50 (right) wo mito-roGFP2-Orp1/WT and mito-roGFP2-Orp1/*Taz*-KD mice. DTT was used as a reductive control (blue). N (WT, 20 wo) = 7, N (*Taz*-KD, 20 wo) = 7, N (DTT, 20 wo) = 8, N (WT, 50 wo) = 4, N (*Taz*-KD, 50 wo) = 4, N (DTT, 50 wo) = 6. (**E**) Real-time H_2_O_2_ imaging of *ex vivo* 20 wo mito-roGFP2-Orp1/WT and mito-roGFP2-Orp1/*Taz*-KD pancreatic islets. Pancreatic islets were incubated with RPMI 1640 at 5% CO_2_ and 37 °C, in presence or absence of 1 µM Auranofin and 100 µM exogenously added H_2_O_2_, N = 5. A schematic figure illustrates the CD7 redox measurement with isolated islets, created with Biorender.com. (**F**) Representative western blot and quantification of Prx3 (left), Cat (middle) and GPX4 (right) normalized to β-actin in pancreatic islets of 20 wo *Taz*-KD mice, N (Prx3) = 4, N (Cat) = 3, N (GPX4) = 4. Data represent mean ± SEM (indicated by error bars); N and n numbers indicate number of animals and experiments, respectively; statistical significance was determined by unpaired Student *t* test: *p < 0.05, **p < 0.01. Abbreviations: Mitochondria-redox-sensitive- GFP2-Orp1 (Mito-roGFP2-Orp1), doxycycline (doxy), weeks of age (wo), *Tafazzin*-Knockdown (*Taz*-KD), Wildtype (WT), Dithiothreitol (DTT), glucose (Glu, G), catalase (Cat), peroxiredoxin 3 (Prx3), glutathionperoxidase 4 (GPX4), area under the curve (AUC).

Subsequently, we isolated pancreatic islets from 20 wo mice and followed the mito-roGFP2- Orp1 oxidation under culture conditions (10 mM of glucose) or in the presence of H_2_O_2_ or the thioredoxin reductase inhibitor, auranofin (Fig. 4E). We saw no difference in steady state probe oxidation or response dynamics in any condition (Fig. 4E). Finally, analysis of different redox proteins showed that while Prx3 protein levels were increased in *Taz*-KD mice, catalase levels were decreased (Fig. 4F, Fig. Suppl. 4C). GPX4, NOX4 and Nrf2 protein levels were unchanged among the genotypes (Fig. 4F, Fig. Suppl. 4B, C).

In summary, we observed that *Taz*-KD mice islets exhibit decreased mito-roGFP2-Orp1 oxidation *in vivo*, and similar response *ex vivo* to exogenous oxidants.

### Defective mitochondrial dynamics and increased mitochondrial volume in pancreatic islets of *Taz*-KD

*Taz*-KD has been linked to decreased mitophagy in mouse embryonic fibroblasts and cardiac tissues (44–46). Furthermore, mitochondrial network morphology is known to dynamically change in response to stress conditions to ensure function and survival (47, 48). Therefore, we were particularly interested to analyze mitochondrial volume and dynamics in *Taz*-KD. Using confocal laser microscopy, we visualized the mitochondrial network and nucleus of pancreatic islet cells with Mitotracker and Hoechst, respectively (Fig. 5A). We observed a significantly increased Mitotracker intensity of the *Taz*-KD dispersed pancreatic islet cells compared to WT, supporting an increased mitochondrial volume (Fig. 5B). Consistent with this observation, a citrate synthase activity assay revealed increased activity in *Taz*-KD pancreatic islet cells (Fig. 5C).

**Figure 5:**
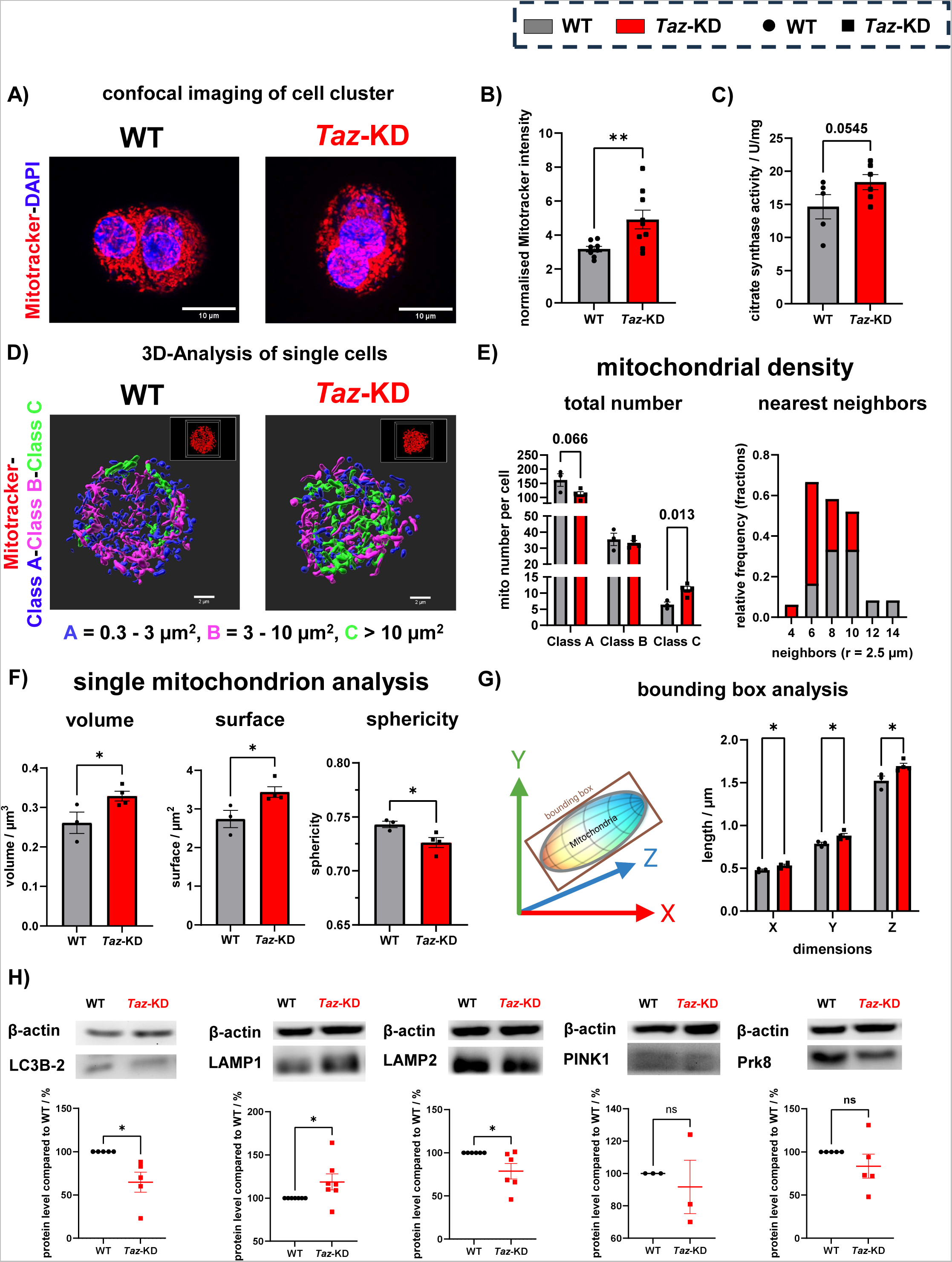
Increased single mitochondrial dimensions lead to overall increase in mitochondrial volume but decrease in number. (**A**) Representative images of dispersed pancreatic islet cell cluster of 20 wo WT and *Taz*-KD mice and mitochondrial volume quantification (**B**) using confocal microscopy with MitoTracker ^TM^ Deep Red and DAPI. Scale bar: 10 µm. N (WT) = 8, N (*Taz*-KD) = 9. (**C**) Citrate synthase activity assay of 20 wo WT and *Taz*-KD pancreatic islets to determine mitochondrial mass normalized to protein content, N (WT) = 5, N (*Taz*-KD) = 6. (**D**) Representative 3D-rendering images of the mitochondrial network from single dispersed pancreatic islet cells of 20 wo WT and *Taz*-KD mice, classified according to their surface area into three different classes: A = 0.3 – 3 µm^2^ (blue), B = 3 – 10 µm^2^ (magenta), C > 10 µm^2^ (green). The respective confocal microscopy image is shown in the upper right. Scale bar: 2 µm. (**E**) Mitochondrial density analysis, including the mitochondrial number (left) per pancreatic islet cell in separate classes (Class A, Class B and Class C) and a frequency distribution histogram (right) using a nearest neighbor analysis testing for neighboring mitochondria in a radius of 2.5 µm from WT and *Taz*- KD dispersed pancreatic islet cells. The frequency distribution histogram displays the fractions of WT and *Taz*-KD mitochondria that have a certain number of neighbors. N (WT) = 3, N (*Taz*- KD) = 4. (**F**) Single mitochondrion analysis of volume (left), surface area (middle), sphericity (right) from 20 wo WT and *Taz*-KD dispersed pancreatic islet cells, N (WT) = 3, N (*Taz*-KD) = 4. (**G**) Schematic figure (left) of object-oriented bounding box analysis in 3D (x, y and z) and quantification (right) of bounding box in single mitochondria from 20 wo WT and *Taz*-KD dispersed pancreatic islet cells. (**H**) Representative western blot and quantification of LC3B-2 (left), LAMP1 (2^nd^ left), LAMP2 (3^rd^ left), PINK1 (4^th^ left) and Prk8 (right) normalized to β-actin in pancreatic islets of 20 wo *Taz*-KD mice, N (LC3B-2) = 5, N (LAMP1) = 7, N (LAMP2) = 6, N (PINK1) = 3, N (Prk8) = 5. Data represent mean ± SEM (indicated by error bars); N numbers indicate number of animals; statistical significance was determined by unpaired or paired (western blot) Student *t* test: *p < 0.05, **p < 0.01. Abbreviations: weeks of age (wo), *Tafazzin*- Knockdown (*Taz*-KD), Wildtype (WT), lysosomal-associated membrane protein 1 (LAMP1), lysosomal-associated membrane protein 2 (LAMP2), parkin (Prk8).

STED microscopy was used to study the mitochondrial network morphology of pancreatic islet cells in greater detail. STED microscopy resolves mitochondrial ultrastructure with a greater detail and sharpness, allowing for a more accurate determination of mitochondrial volume. As expected, STED microscopy reported a 38.75 ± 3.86% decrease in total mitochondrial volume and a 30.69 ± 11.24% increase in mitochondrial number compared to that suggested by confocal microscopy (Fig. Suppl. 5A, B). After imaging and 3D-rendering, we classified mitochondria into three different categories based on surface area: small (Class A): 0.3 – 3 µm^2^, medium (Class B): 3 – 10 µm^2^ and large (Class C): > 10 µm^2^. We observed an increase in Class C and decrease in Class A mitochondria in *Taz*-KD pancreatic islet cells (Fig. 5D, E). A nearest neighbor analysis showed a decreased frequency of mitochondrial neighbors in *Taz*- KD islet cells (Fig. 5E), which is consistent with the decrease in overall mitochondrial number due to a more connected mitochondrial network in *Taz*-KD (Fig. Suppl. 5C). Individual mitochondria of *Taz*-KD displayed increased volume and surface area but decreased sphericity (Fig. 5F) due to the shift towards bigger mitochondria, as the single mitochondrion morphology in separated classes remained unchanged (Fig. Suppl. 5D). A bounding box analysis confirmed the enlargement of *Taz*-KD mitochondria, which is not directed, but instead similar on each axis (X, Y and Z) (Fig. 5 G and Fig. Suppl. 5E).

Defective autophagy and mitophagy were previously described in heart and skeletal muscle tissue of mouse *Taz*-KD samples (45, 46). Therefore, we investigated essential autophagy and mitophagy proteins using western blot analysis. We observed decreased protein levels of LC3B2, an essential protein in autophagic initiation that is directly correlated with the number of autophagosomes in *Taz*-KD pancreatic islets (Fig. 5H, Fig. Suppl. 5E). Its cytosolic isoform, LC3B1, was also reduced in *Taz*-KD (Fig. Suppl. 5F). Atg7, a driver of LC3B conjugation tended to be reduced in *Taz*-KD (Fig. Suppl. 5G). The protein level of LAMP1 was increased while its functional analog LAMP2 was decreased in *Taz*-KD pancreatic islets (Fig. 5H, Fig. Suppl. 5H). The protein levels of the mammalian mitophagy proteins PINK1 and Prk8 did not significantly change (Fig. 5H, Fig. Suppl. 5H).

In summary, we observe altered mitochondrial dynamics leading to enlarged and less round single mitochondria, an increased mitochondrial volume, and a decreased overall mitochondrial number in *Taz*-KD pancreatic islets.

### *Taz*-KD leads to changes in genes involved in N-acetyl-glucosamine metabolic process and Protein-*O*-linked glycosylation

To gain deeper insight into different pathways and mechanisms contributing to the changes in pancreatic islets of 20 wo *Taz*-KD mice, we performed whole islet bulk mRNA-sequencing. We observed significant downregulation of 66 genes, while 98 genes were upregulated in *Taz*-KD pancreatic islets compared to WT (Fig. 6A). Heatmaps representing the top 30 up- and down-differentially expressed genes (DEG), (protein coding) are shown for different WT and *Taz*-KD samples (Fig. 6B). Gene ontology (GO) terms were analyzed and selected significantly downregulated DEG included purine nucleotide metabolic and biosynthetic processes *(Apoe, Taz, mt-ND4l*) and regulation of immune response (*Cd74, H2-Ab1, H2-Aa*) (Fig. 6C, top panel). Increased GO terms included DEG involved in N-acetyl-glucosamine metabolic process (*Gnpnat1*) and Protein-O-linked glycosylation (*B3gnt9, Galnt17, Gcnt7*) (Fig. 6C, bottom panel, Fig. Suppl. 6A). Reactome pathway analysis showed enrichment in “Protein-O-linked glycosylation” and “*O*-linked β-*N*-acetylglucosamine” (*O*-GlcNAc) pathways, summarized in the schematic figure, where significantly upregulated genes in *Taz*-KD are shown in purple (Fig. 6D). Due to the particular importance of these pathways in nutrient sensing and metabolism of pancreatic islets (49), we further investigated *O*-GlcNAc protein modification in pancreatic islets. Using an antibody against *O*-GlcNAc, we found enhanced levels of *O*-GlcNAc protein modification by immunohistochemistry (Fig. 6E) and western blot (Fig. 6F and Fig. Suppl. 6B) in 20 wo *Taz-*KD pancreatic islets compared to WT.

**Figure 6:**
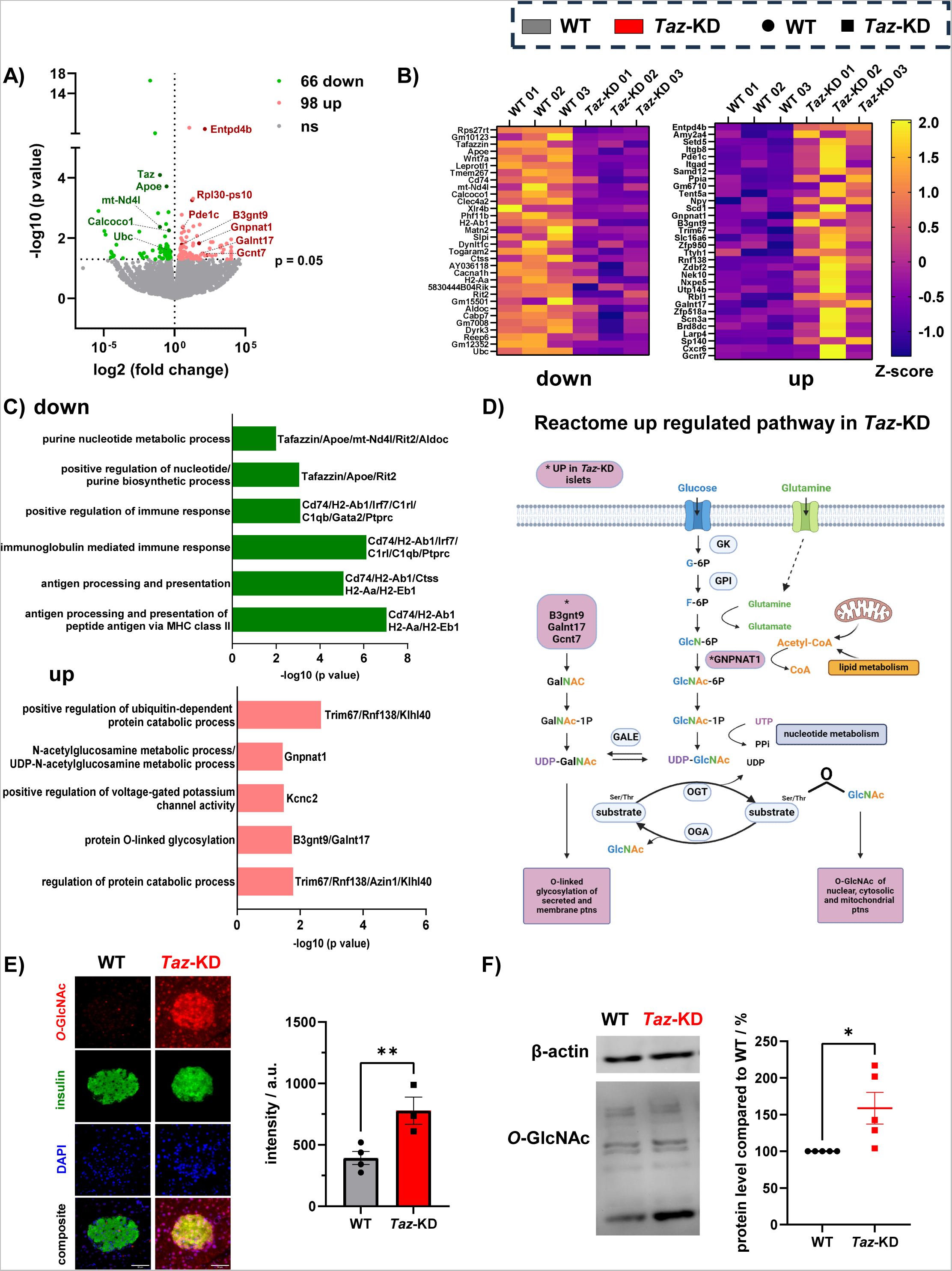
RNA-Sequencing reveals upregulation of *O*-GlcNAc in *Taz*-KD pancreatic islets. Analysis of pancreatic islet mRNA sequencing from 20 wo WT and *Taz*-KD mice are shown in **A** to **C**, N = 3. (**A**) Volcano plot illustrates the upregulation of 98 genes and downregulation of 66 genes (p > 0.05) in *Taz*-KD. Selected genes that are significantly altered in *Taz*-KD pancreatic islets are highlighted, N = 3. (**B**) Heatmap of the 30 most down- (left panel) and upregulated (right panel) genes normalized to the Z-score. (**C**) Identified relevant GO pathways with genes that are down- (left panel) or upregulated (right panel) in *Taz*-KD. (**D**) Schematic figure of the *O*-GlcNAc pathway, upregulated in Reactome analysis of *Taz*-KD. The proteins and pathways resulting from the upregulated genes in *Taz*-KD are highlighted in purple. (**E**) Representative IHC images (left) and intensity quantification (right) of 20 wo WT and *Taz*-KD pancreatic islets stained against *O*-GlcNAc (red), insulin (green), and DAPI (blue). N (WT) = 4, N (*Taz*-KD) = 3. (**F**) Representative western blot (left) and quantification (right) of *O*-GlcNAc normalized to β-actin in pancreatic islets of 20 wo *Taz*-KD mice, N = 5. Data represent mean ± SEM (indicated by error bars); N numbers indicate number of animals; statistical significance was determined by unpaired or paired (western blot) Student *t* test: *p < 0.05, **p < 0.01. Abbreviations: *O*-linked β-*N*-acetylglucosamine (*O*-GlcNAc), weeks of age (wo), *Tafazzin*- Knockdown (*Taz*-KD), Wildtype (WT), gene ontology (GO), immunohistochemistry (IHC).

In summary, bulk mRNA-sequencing revealed alterations in genes involved in metabolism, immune response, and nutrient sensing in *Taz*-KD pancreatic islets.

### *In vivo Taz*-KD leads to increased glucose tolerance and with preserved insulin and glucagon secretion

Finally, we investigated if the alterations observed in *Taz*-KD pancreatic islets metabolic pathways would impact whole-body glucose homeostasis. During an intraperitoneal glucose tolerance test (i.p. GTT) (Fig. 7A), *Taz*-KD mice at 20 and 50 wo mice displayed significantly reduced blood glucose levels 60 and 120 min after glucose injection when compared to the respective WT group (Fig. 7B, Fig. Suppl. 7A, B). Additionally, plasma insulin and glucagon levels were measured during i.p. GTT. Upon glucose injection, *Taz*-KD plasma insulin levels reached similar values as WT, which was also observed when fold increase was calculated (Fig. 7C, Fig. Suppl. 7C), suggesting a preserved response to glucose. Interestingly, in *Taz*-KD mice, the plasma glucagon levels show a different profile, reaching a peak at 15 min, while WT mice show a decrease in glucagon levels in the first minutes, albeit not statistically significant (Fig. 7D, Fig. Suppl. 7D).

**Figure 7:**
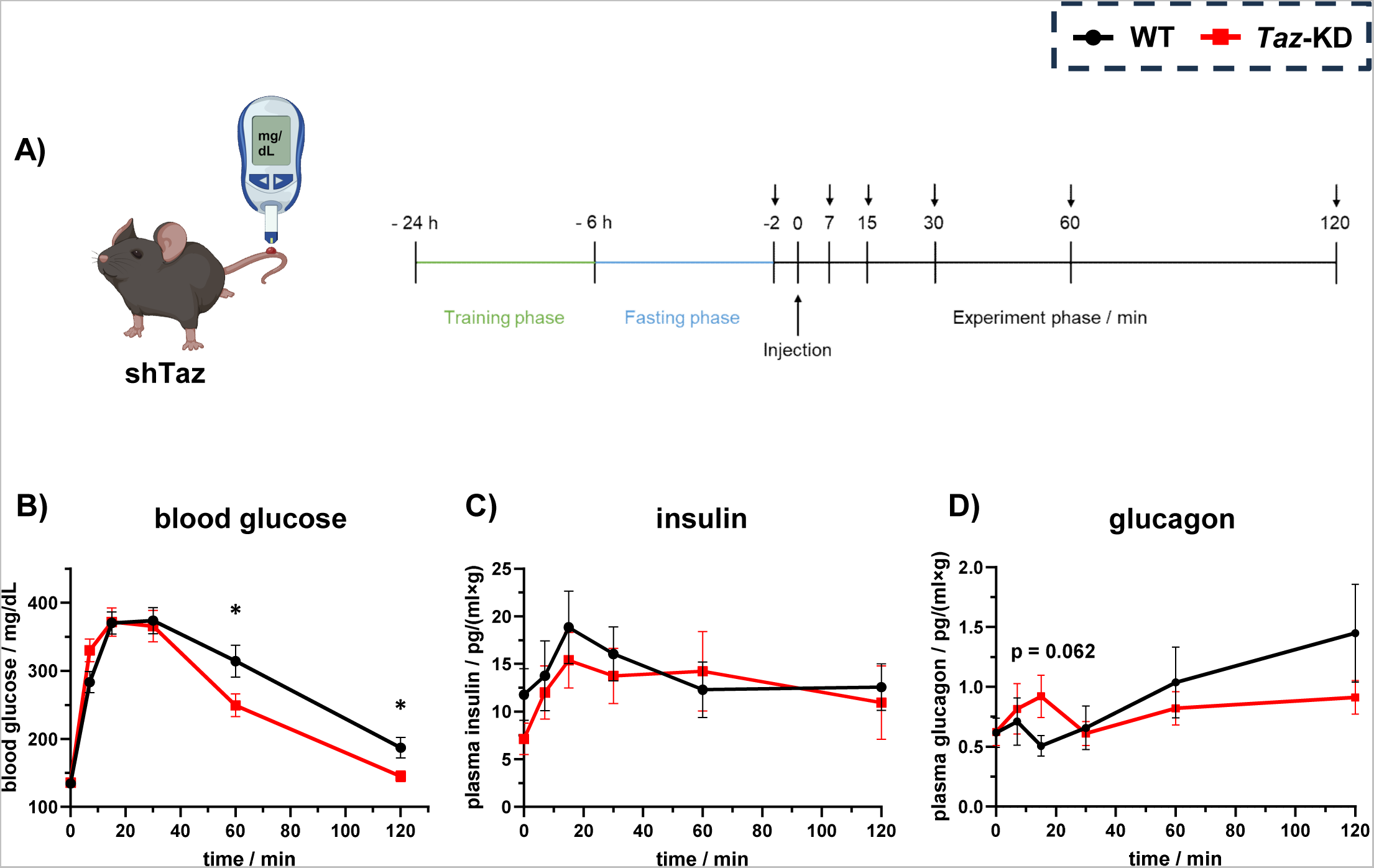
Alterations in *Taz*-KD whole-body glucose homeostasis in GTT. (**A**) Schematic illustration of blood glucose measurement (left) and protocol of the GTT (right) of shTaz mice, which is performed after an initial training (24 h before GTT) and fasting (6 h before GTT) phase. The arrows above the time line indicate measurement during the experiment phase. Illustration created with Biorender.com. GTT of *Taz*-KD and WT mice at 20 wo with measured blood glucose (**B**), plasma insulin (**C**) and plasma glucagon (**D**) levels, N (blood glucose levels, WT) = 24, N (blood glucose levels, *Taz*-KD) = 22; N (plasma insulin and glucagon, WT) = 7, N (plasma insulin and glucagon, *Taz*-KD) = 8. Data represent mean ± SEM (indicated by error bars); N numbers indicate number of animals; statistical significance was determined by unpaired Student *t* test: *p < 0.05. Abbreviations: weeks of age (wo), *Tafazzin*- Knockdown (*Taz*-KD), Wildtype (WT), *Tafazzin* (*Taz*), glucose tolerance test (GTT).

In summary, we show that *in vivo Taz*-KD for 20 weeks leads to decreased CL levels and changes in lipid profile in pancreatic islets. However, pancreatic islets undergo a series of adaptations including cellular compositional changes as well as transcriptional, mitochondrial morphological and metabolic changes which preserve islet function and maintain *in vivo* insulin levels. Redox changes seem to play a minor role. Interestingly, increased glucagon secretion and α-cell number is also observed in this model (Fig 8A).

**Figure 8:**
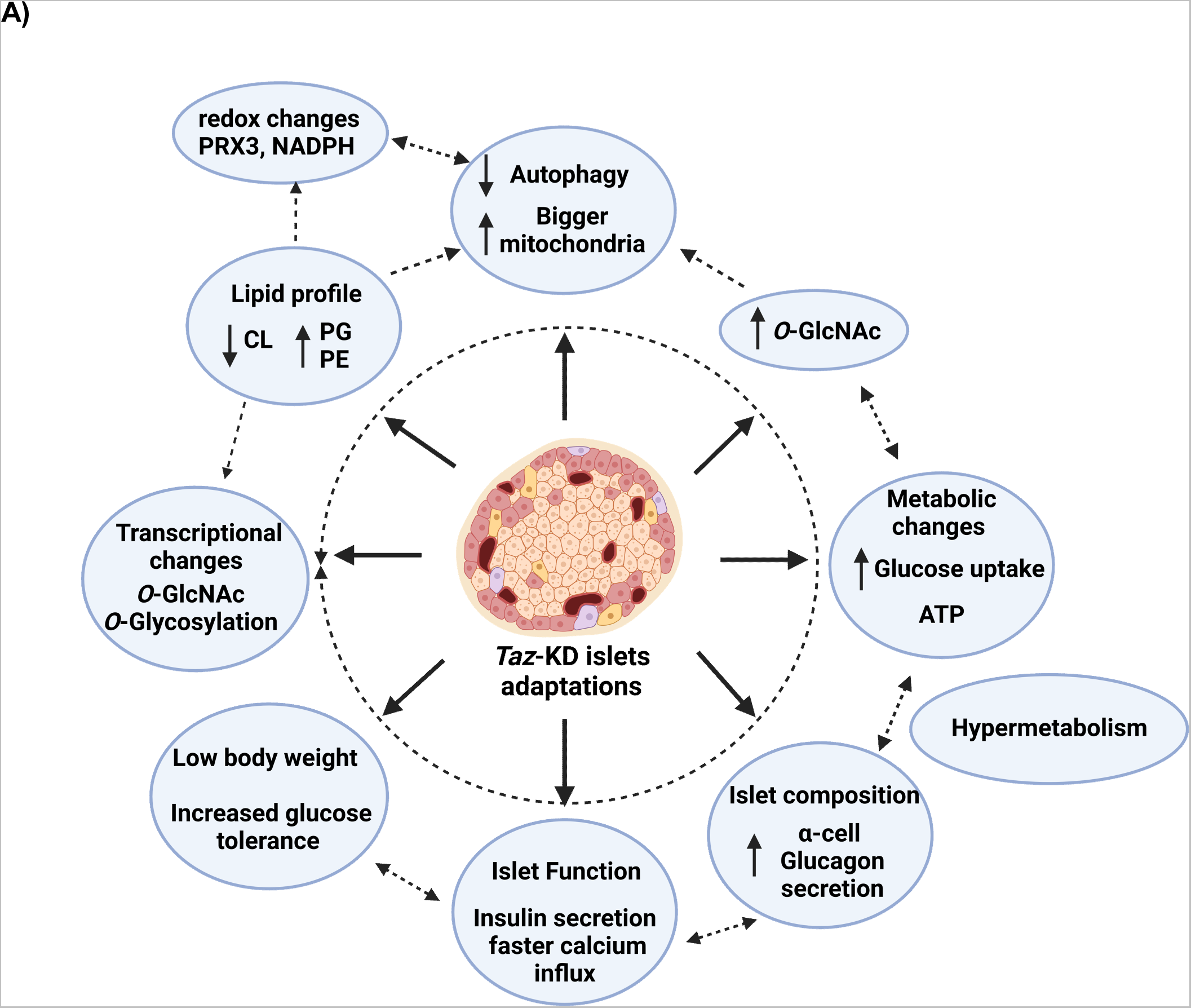
Adaptive processes which preserve pancreatic islet function in *Taz*-KD. (**A**) Summary of adaptative processes on pancreatic islet biology as a consequence of *Taz*- KD observed in a shTaz strain at 20 wo. Abbreviations: weeks of age (wo), *Tafazzin*- Knockdown (*Taz*-KD), Wildtype (WT), *Tafazzin* (*Taz*), *O*-linked β-*N*-acetylglucosamine (*O*- GlcNAc), cardiolipin (CL).

## Discussion

Deficiency in secretion of pancreatic islet hormones (insulin, glucagon, and somatostatin) leads to metabolic imbalances such as insulin resistance, diabetes, and related co-morbidities. Pancreatic islet adaptation is essential for maintaining whole-body glucose homeostasis in diverse situations including pregnancy, aging, inflammation, and metabolic diseases. Here, we show that in a mouse model of BTHS, pancreatic islets undergo a series of metabolic and morphological changes, including increase in mitochondrial volume and alterations in islet cell composition, while maintaining GSIS and upregulating glucagon secretion. Transcriptomic analyses revealed increased expression of genes encoding enzymes involved in GlcNAc synthesis, whilst western blot analysis showed increased protein *O*-linked GlcNAc modification. This potentially serves as a link between glucose metabolism and mitochondrial adaptations through regulation of autophagy (50–52).

Mutations in the *Taz* gene lead to development of BTHS due to the decreased CL remodeling. BTHS primarily affects heart function, however, endocrine and metabolic effects have also been described in patients, including recurrent hypoglycemia, hypocholesterolemia, delayed puberty and 3-methylglutaconic aciduria (53). Surprisingly, despite the importance of CL for cellular function, the effects of CL loss or changes in CL remodeling were rarely addressed in pancreatic islets (54, 55), especially with regard to the role of the *Taz* acyltransferase which, to the best of our knowledge, has been addressed only once before (21). In pancreatic islets, *Taz*-KD leads to a decrease in CL levels and changes in relative abundance of CL species and other phospholipids. Similar to findings in human patients (56), our data show that in a mouse model of BTHS, *Taz*-KD leads to improved glucose tolerance, which we show is not due to increased plasma insulin levels, and likely reflects increased insulin sensitivity in peripheral tissues such as heart, skeletal muscle, lymphocytes (9, 14). *Taz*-KD pancreatic islets also showed increased glucose uptake and NAD(P)H levels, with no changes in ATP production and whole islet respiration. The use of different metabolic substrates and the adaptive mechanisms involved in meeting energy demands in BTHS are poorly understood. We speculate that increased glucose uptake is an adaptation to early inefficient mitochondrial metabolism. Interestingly, hypermetabolism is often associated with mitochondrial disorders (57). In addition, increase in mitochondrial volume has also been shown to be a successful strategy to maintain respiration, decrease ROS accumulation and promote cell survival during stress and cell proliferation (57–59).

Oxidative stress and increased ROS production have been suggested as central players in the development of the BTHS phenotype. However, using a redox histology approach, we observed decreased mitochondrial H_2_O_2_ levels in 20 and 50 wo *Taz*-KD mice. Similar observations were also reported in the heart of *Taz*-KD mice (8, 39). These results are in contrast to other studies that have reported increased oxidative stress in tissues from BTHS mice and patient samples (60, 61). The reason for these differences probably relies on the methodology used (*in vivo* versus *ex vivo,* incubation protocols), as the use of chemical dyes that lack redox species specificity exhibit poor subcellular targeting, and are prone to a range of artefacts (62, 63). In our study, reduced H_2_O_2_ levels could be a result of lower H_2_O_2_ production, increased scavenging capacity (Prx3) or increased availability of reducing equivalents, such as NADPH. Indeed, in *Taz*-KD mice, NAD(P)H levels were increased and can be used as co-factor by antioxidant enzymes such as GSH/GRX, PRDX and TRX.

Interestingly, *Taz*-KD leads to changes in islet cell composition, resulting in increased α-cell number. The altered islet composition in the *Taz*-KD mice resulted in increased glucagon secretion in low glucose, without any changes in glucose-stimulated insulin secretion. The reason for increased α-cell number is unclear. Interestingly, in mouse models of mitochondria dysfunction, where complex I expression was decreased, pancreatic α-cell mass was also increased (64). Whether increase in α-cells is due to neogenesis or transdifferentiating cells is also unknown, but we did not observe any significant differences in markers for proliferation or cell death. β- to α-cell transdifferentiation were observed in diverse models of diabetes and insulin resistance (65) and could also play a role in this model. Increased α-cell number was also observed in the only other study on pancreatic response to *Taz* knockdown (21). Taken together, it would be extremely interesting to look for differences in α-cell number and the role of glucagon in human BTHS patients.

*Taz*-KD islets show faster calcium influx when stimulated by high glucose suggesting that these channels could be in a state of pre-activation. Due to the fact that 1) *Taz*-KD islets have more α-cells and higher glucagon secretion, 2) it has been shown that glucagon secretion act on the glucose set-point of islets (66) and 3) activation of glucagon receptor (and potentially GLP-1 receptor) (67), may lead to intracellular calcium mobilization, (similar to GLP-1 receptor activation) (68, 69), it is tempting to speculate that increased glucagon secretion and hence, amplified paracrine signaling could also be responsible for the effects on calcium influx in pancreatic β-cells (70). Supporting this idea, cytosolic calcium influx was similar in both genotypes when pancreatic islets were dispersed into isolated cells therefore removing the close α-to-β cell contact.

Excessive mitochondrial fusion or fission has been implicated in pancreatic β-cell gain or loss of function (71, 72). CL serves as a signaling platform and was shown to be important for proper mitochondrial function and dynamics (reviewed in (73)). Here, we show for the first time in pancreatic islets, that CL remodeling is important for regulation of mitochondrial dynamics, as demonstrated by increased mitochondrial volume in *Taz*-KD islets. In line with other studies in different BTHS cell types (45, 46), we observed that *Taz*-KD leads to changes in autophagy and mitophagy markers, which may be responsible changes in mitochondrial network dynamics. During mitochondrial stress or damage, CL can be externalized to the outer mitochondrial membrane, acting as a signal for the recruitment of mitophagy-related proteins, including LC3, a protein involved in autophagosome formation. Here, we showed that LC3B was decreased in pancreatic islets of *Taz*-KD mice. Mitochondria morphology and dynamics was shown to depend on the interplay of several lipids (74). We also show an increase in PG and PC in *Taz*-KD islets, which have previously been related to changes in mitochondrial morphology, including in other BTHS models (74–76).

*In vivo* RNAseq analysis showed that *O*-GlcNAcylation and *O*-glycosylation pathways were increased in *Taz*-KD pancreatic islets, which was further confirmed for *O*-GlcNAcylation by western blot and immunohistochemistry. *O*-GlcNAcylation is a post-translational modification where *O*-GlcNAc transferase (OGT) adds a N-acetylglucosamine (GlcNAc) sugar moiety to serine or threonine residues on nuclear and cytoplasmic proteins. This modification is dynamic and reversible, and is involved in various cellular processes, including signaling, transcription, and metabolism, and has emerged as a crucial regulator of mitochondrial function and quality control mechanisms, including mitophagy (77). *O*-GlcNAcylation is recognized as one of the essential regulators of cell signaling, equivalent to the well-studied phosphorylation (78). *O*- GlcNAcylation is particularly important for pancreatic β-cells mass and function, as they uniquely express high levels of OGT (79, 80) and are optimized to sense nutrient flux. Furthermore, *O*-GlcNAcylation was also shown to be involved in β-cell adaptation during compensatory pre-diabetic phase (81). Moreover, increased *O*-GlcNAcylation was shown to blunt autophagic signaling in the diabetic heart (51), resulting in elongated mitochondria (52). In a pancreatic β-cell line, Min6, increased *O*-GlcNAcylation resulted in preserved GSIS during chronic hyperglycemia, possibly through increases in mRNA levels of *Ins1* and *Ins2* via epigenetic modifications (82). Our RNAseq data also showed that several chromatin modulators and regulation of transcription and translation (Brd8dc, Zfp518a, Larp4, Sp140, RNF138, Zfp950) are increased in *Taz*-KD pancreatic islets. Further experiments are necessary to understand the molecular targets and consequences of increased *O*- GlcNAcylation in pancreatic islets of *Taz*-KD mice.

Of note, circulating factors could also play a role in islet adaption in the context of BTHS. For example, it was recently shown that the growth differentiation factor 15 (GDF15) was elevated in BTHS patients (83). GDF15 exerts a positive role in pancreatic β-cell insulin secretion, glucose uptake (84, 85), and improve pancreatic β-cell survival and inflammation (86). Whether GDF15 is also involved in pancreatic islets adaptation in *Taz*-KD mouse is unknown.

In summary, in a mouse model of Barth Syndrome, we have demonstrated that pancreatic islet function is robustly maintained. Pancreatic islets exhibit cellular compositional, morphological, transcriptional and metabolic changes that are likely important mechanisms allowing the adaptation towards decreased CL remodeling. Our data provide new insights into the metabolic effects associated with BTHS, and additionally, on the role of CL remodeling for pancreatic islet function and metabolic diseases. With recent advances in the management of heart failure in BTHS patients (main cause of death), and prolonged life expectancy, gaining knowledge regarding secondary effects of BTHS in less affected tissues is of great importance, especially as the BTHS patients get older and might be challenged with different diets and metabolic complications. Pancreatic islets play a central role in coordinating whole-body energy and metabolic status and understanding the role of CL remodeling and BTHS phenotype on pancreatic islets function will also bring new insights into the diabetes field.

## Acknowledgements

We thank Angélique Schniebs, Andrea Armbrüster, Sandra Janku and Ruth Nickels for excellent technical assistance. BM is grateful for generous support from the German Research Foundation (DFG), MO 2774/7-1 project number: 508372800, and MO 2774/6-1 project number 505680640. We also thank Gebhard Stopper for software and data analysis support. CC was supported by the SFB894. LPR is grateful for support from DFG SFB894 and TRR 219 Project ID 322900939.

## Data availability

The complete gene list of RNAseq data and lipidomics profile information (including lipid class, species, double bond, hydroxylation, carbon length and fatty acid profiles) will be made available upon request.

## Supplementary information

### PCR protocols

**ShTaz** 95 °C 5 min – (95 °C 30 sec – 60 °C 30 sec – 72 °C 1 min) x40 – 72 °C 10 min

**Orp** 95 °C 3 min – (95 °C 30 sec – 64 °C 30 sec – 72 °C 1 min) x20 – (95 °C 30 sec – 54 °C 30 sec – 72 °C 1 min) x10 – 72 °C 5 min

### Primer sequences

Rosa JD 72: 5’ CCA TGG AAT TCG AAC GCT GAC GTC 3’

Rosa JD 73: 5’ TAT GGG CTA TGA ACT AAT GAC CC 3’

Rosa JD 74: 5’ GAG ACT CTG GCT ACT CAT CC 3’

Rosa JD 75: 5’ CCT TCA GCA AGA GCT GGG GAC 3’

MS 282: 5’ AAA GTC GCT CTG AGT TGT TAT 3’

MS 284: 5’ GGA GCG GGA GAA ATG GAT ATG 3’

MS 305: 5’ GGG CTA TGA ACT AAT GAC CCC G 3’

E8 for Bl/6N: 5’ TAT TGG CTA CAC AGA CCT TCC 3’

E8_rev for Bl/6N: 5’ TGA CGT GAC TCA TTG TAC CA 3’

E6-12 for Bl/6J: 5’ GTA GGG CCA ACT GTT TCT GC 3’

E6-12_rev for Bl/6J: 5’ TCC CCT CCC TTC CAT TTA GT 3’

### Antibody list

**Supplementary Table 1:**
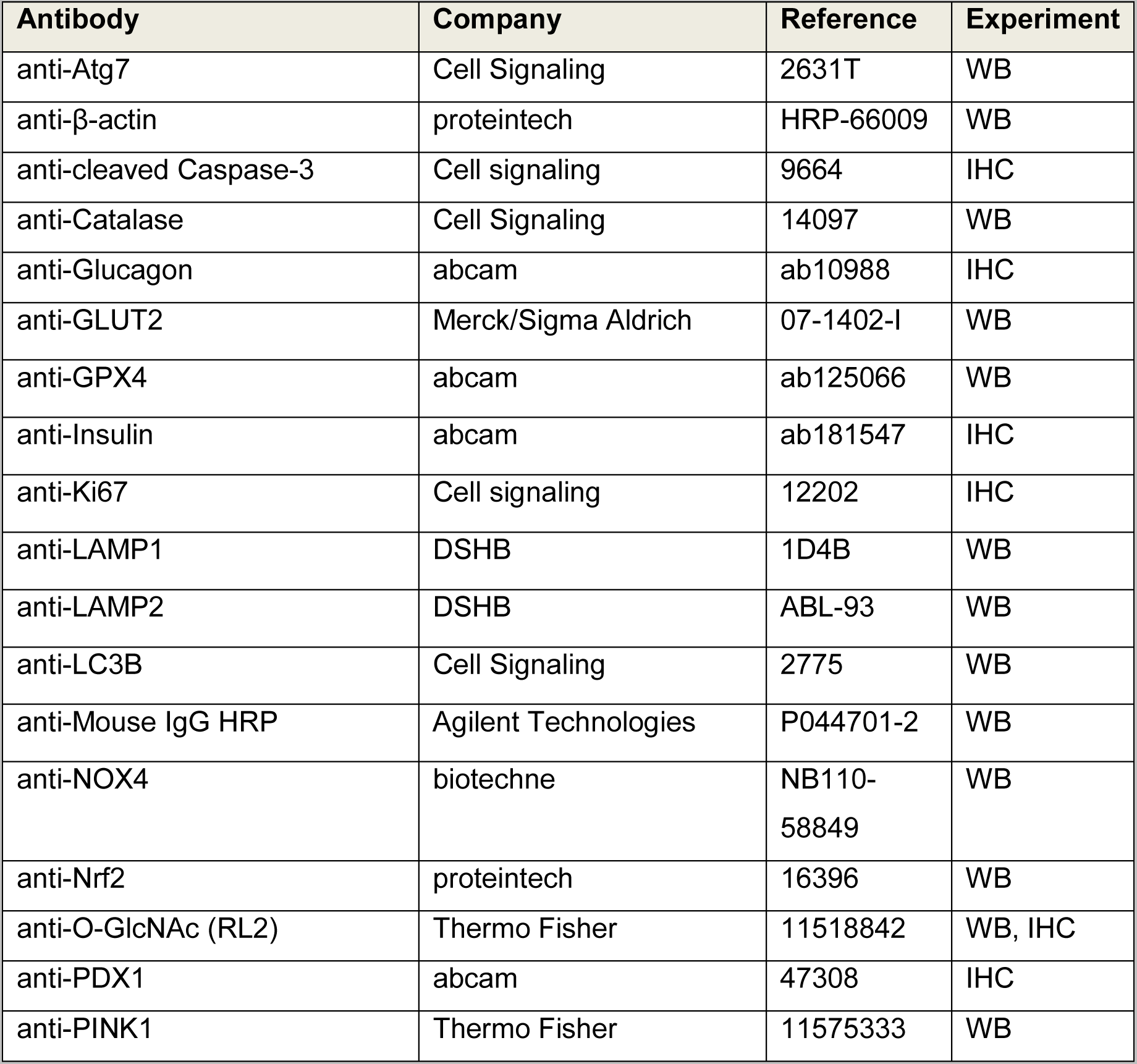
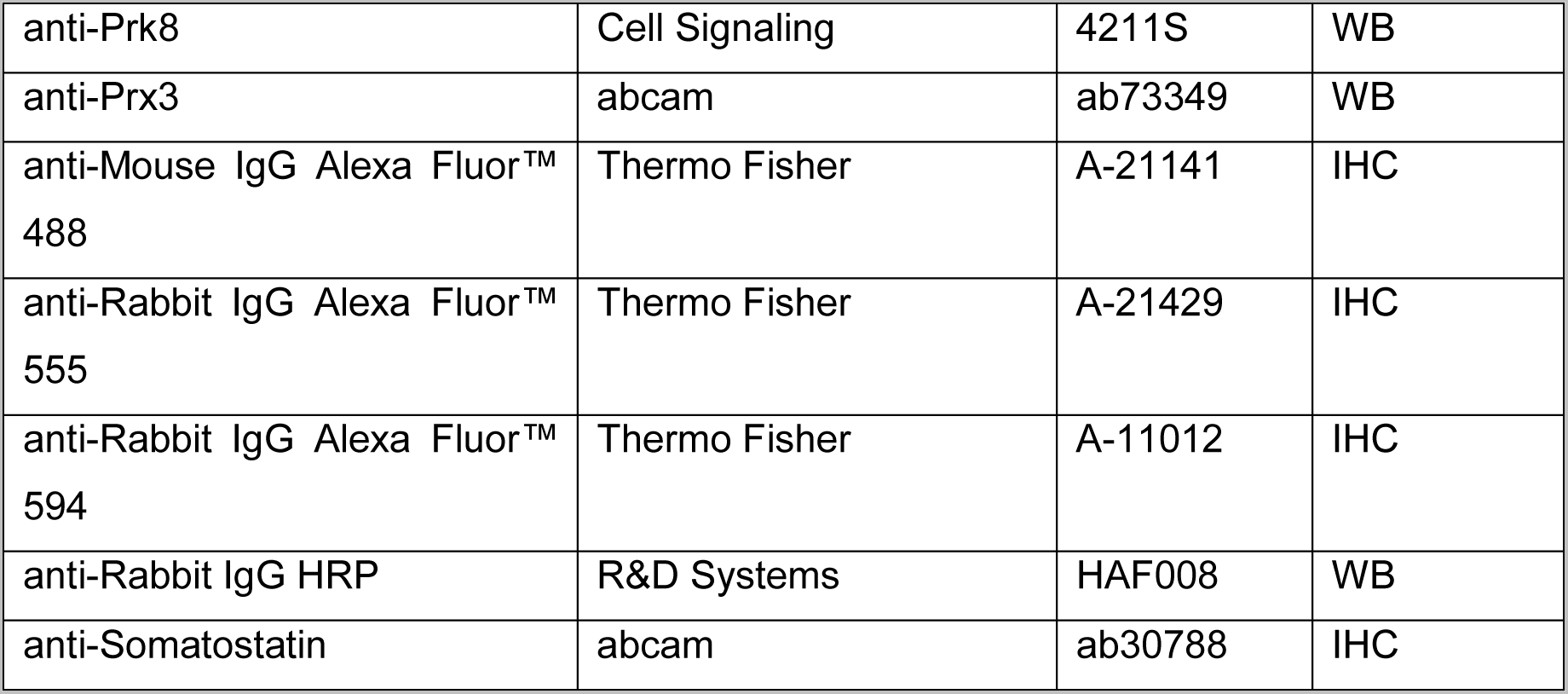
Antibodies used for WB or IHC.

**Supplementary Table 2:** QPCR TaqMan primers.

| Primer | company | Assay ID |
| --- | --- | --- |
| Tafazzin | Thermo Fisher | Mm00504978_m1 |
| GAPDH | Thermo Fisher | Mm99999915_g1 |

**Supplementary Table 3:** Microscopes, plate reader, objectives, and accessories. Abbreviations: numerical aperture (NA)

| device | Company, country | NA |
| --- | --- | --- |
| Axio Observer 7 microscope | Zeiss, Germany |  |
| Axio Observer air objective 20x | Zeiss, Germany | 0.75 |
| Axio Observer oil objective 63x | Zeiss, Germany | 1.4 |
| Celldiscoverer 7 microscope | Zeiss, Germany |  |
| Celldiscoverer 7 air objective 20x | Zeiss, Germany | 0.95 |
| Celldiscoverer 7 water objective 50x | Zeiss, Germany | 1.2 |
| Clariostar plate reader | BMG, Germany |  |
| Stereo microscope 305 | Zeiss, Germany |  |
| Stereo microscope AxioCam 105 color | Zeiss, Germany |  |
| STED microscope | Olympus, Germany |  |
| STED microscope oil objective 100x | Olympus, Germany | 1.4 |

## Additional information

All substances, if not indicated differently were purchased by Sigma/Merck, VWR, Thermo Fisher, Roth and Biomol. Filtersets and beamsplitter for inverted epifluorescence microscopy were manufactured by Zeiss.

## Abbreviation of Lipids

Cholesterol esters (CE), Ceramide (Cer), Cardiolipin (CL), Diacylglycerol (DAG), Hexosylceramide (HexCer), lyso-Phosphatidate (LPA), lyso-Phosphatidylcholine (-ether) (LPC O-), lyso-Phosphatidylethanolamine (-ether) (LPE O-), lyso-Phosphatidylinositol (LPI), lyso- Phosphatidylserine (LPS), Phosphatidate (PA), Phosphatidylcholine (-ether) (PC O-), Phosphatidylethanolamine (-ether) (PE O-), Phosphatidylglycerol (PG), Phosphatidylinositol (PI), Phosphatidylserine (PS), Sphingomyelin (SM), Triacylglycerol (TAG)

---

**Supplementary Figure 1:**
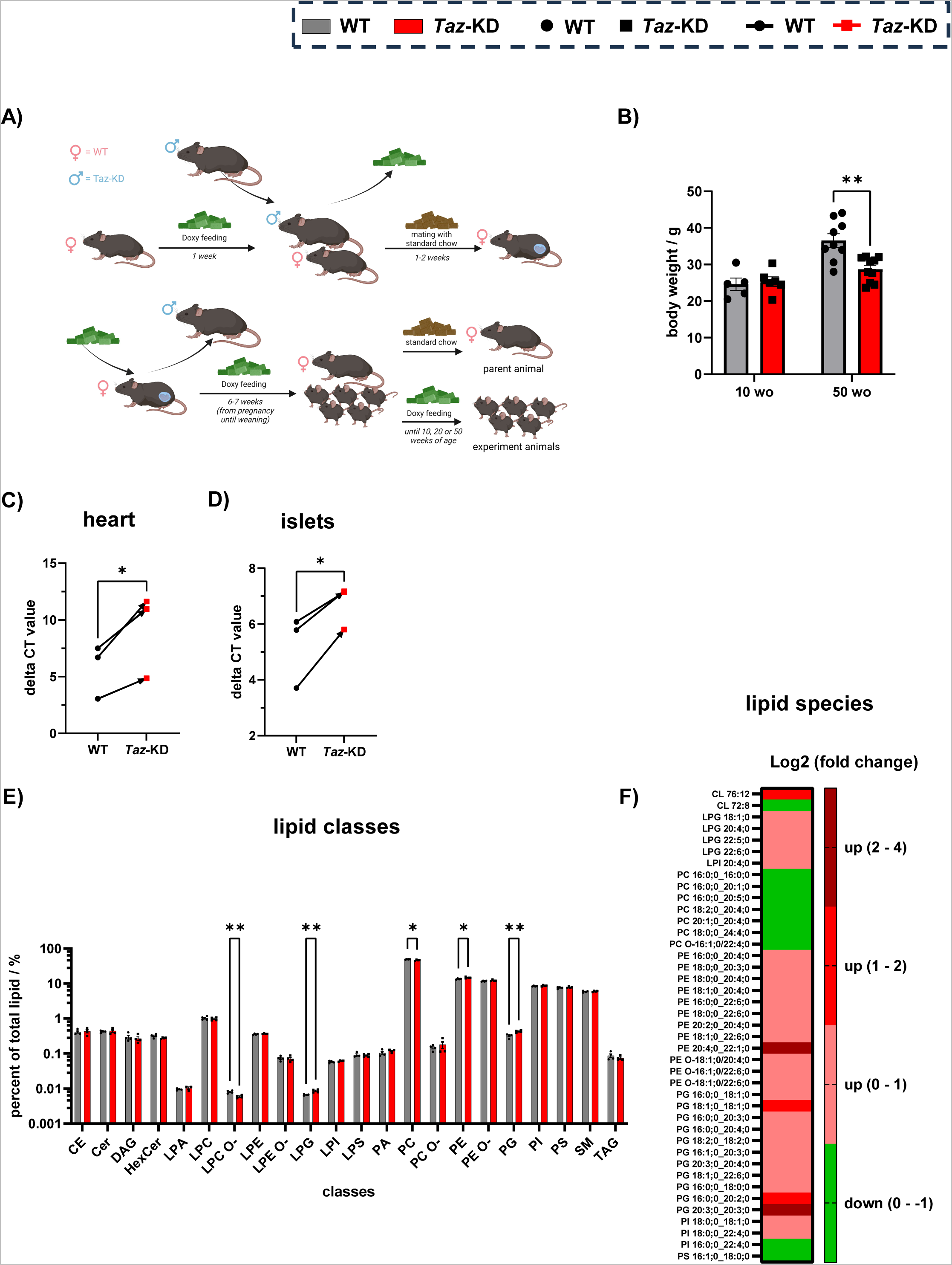
(A) ShTaz doxycycline breeding scheme. (**B**) Body weight of *Taz*-KD and WT at 10 and 50 wo, N (10 wo, WT) = 5, N (10 wo, *Taz*-KD) = 6, N (50 wo, WT) = 9, N (50 wo, *Taz*-KD) = 10. Paired analysis of *Taz* gene expression in heart (**C**) and pancreatic islet (**D**) tissue, N = 3. (**E**) Complete lipid class profile (logarithmic scaling) of pancreatic islets from 20 wo WT and *Taz*- KD, N = 4. (**F**) Significantly (p < 0.05) altered lipid species in *Taz*-KD pancreatic islets. Data represent mean ± SEM (indicated by error bars); N numbers indicate number of animals; statistical significance was determined by unpaired Student *t* test: *p < 0.05, **p < 0.01, ****p < 0.0001. Abbreviations: weeks of age (wo), *Tafazzin*-Knockdown (*Taz*-KD), Wildtype (WT), area under curve (AUC), glucose tolerance test (GTT), lipid class abbreviations can be found in (Supplementary material: Abbreviation of lipids).

**Supplementary Figure 2:**
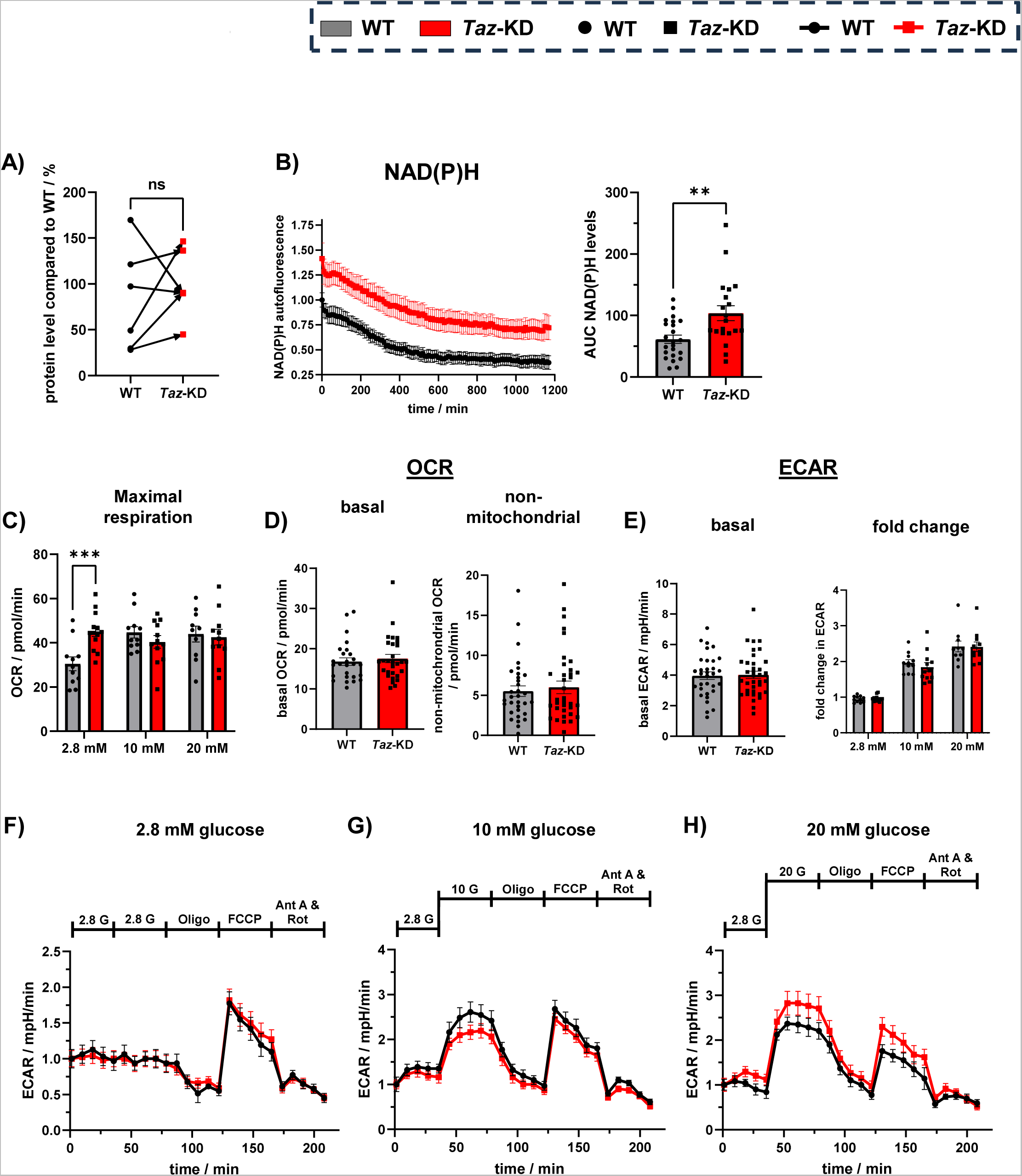
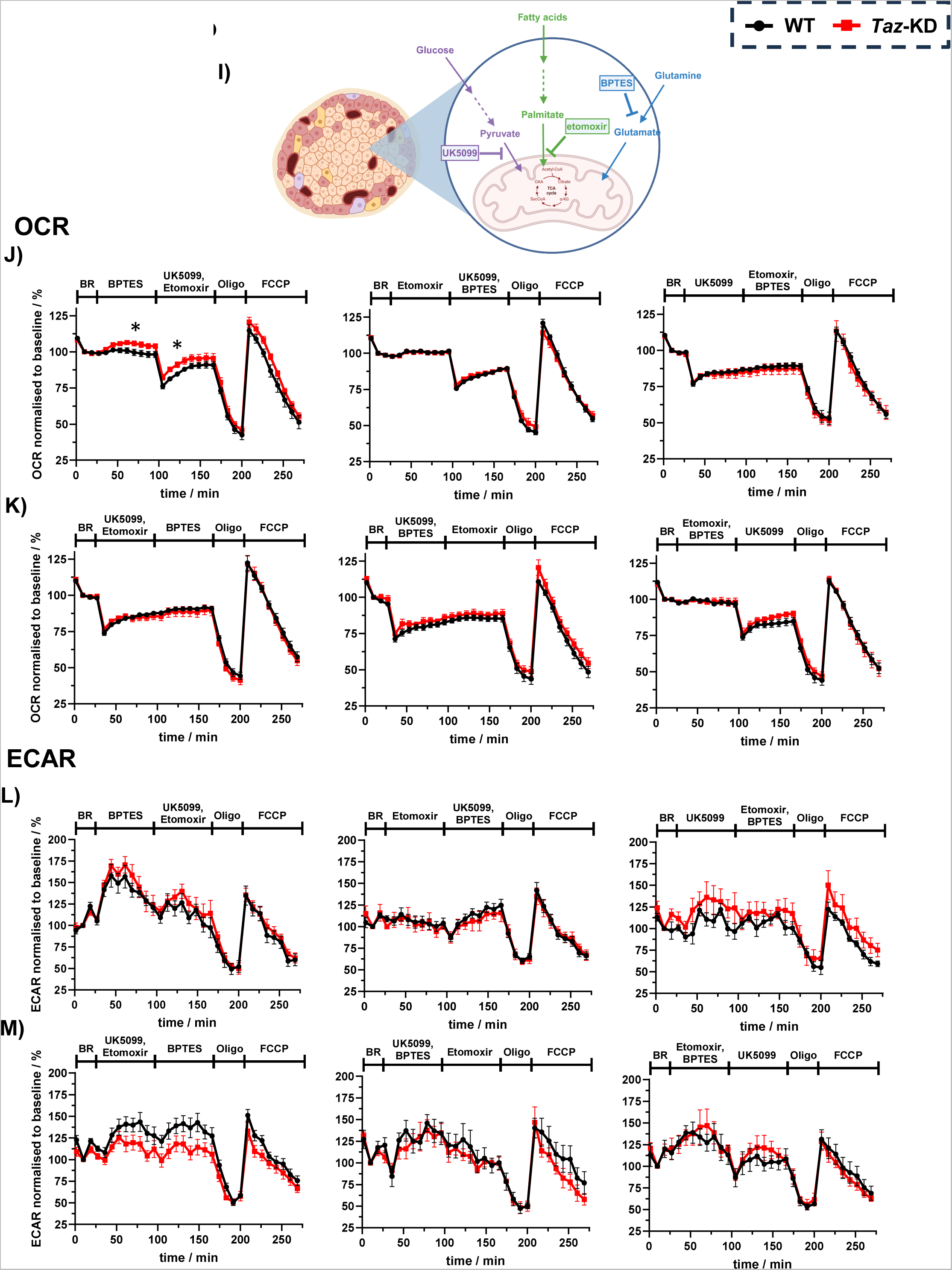
(A) Paired analysis of GLUT2 protein levels in 20 wo WT and *Taz*-KD pancreatic islets, N = 6. (B) NAD(P)H autofluorescence measurement (left) and AUC analysis (right) of 20 wo mito- roGFP2-Orp1/WT and mito-roGFP2-Orp1/*Taz*-KD pancreatic islets in parallel to H_2_O_2_ recordings, n (WT) = 23, n (*Taz*-KD) = 20, from 8 animals. (**C**) Calculated maximal respiration from OCR of WT and *Taz*-KD 20 wo pancreatic islets. n (WT) = 11, n (*Taz*-KD) = 13, n number of experiments include N (WT) = 5 and N (*Taz*-KD) = 4. (**D**) Quantification of basal (left) and non-mitochondrial (right) OCR levels of WT and *Taz*-KD 20 wo pancreatic islets. n (WT) = 32, n (*Taz*-KD) = 36 (**E**) Quantification of basal ECAR (left), and ECAR fold change (right) in response to glucose (2.8, 10, and 20 mM) of WT and *Taz*-KD 20 wo pancreatic islets. n (basal ECAR, WT) = 32, n (basal ECAR, *Taz*-KD) = 36, n (ECAR fold change, WT) = 11, n (ECAR fold change, *Taz*-KD) = 12. ECAR kinetic curves of 20 wo WT and *Taz*-KD pancreatic islets in response to 2.8 mM (**F**), 10 mM (**G**) and 20 mM (**H**) glucose stimulation followed by the addition of inhibitors of the respiratory chain complexes (Oligo, Ant A and Rot) and uncoupler (FCCP). n (WT) = 11, n (*Taz*-KD) = 13. (**I**) Schematic figure of XF Mito Fuel Flex Test Kit protocol. The three inhibitors UK5099, etomoxir, and BPTES are used to inhibit glucose, fatty acid, and glutamine metabolism inside the mitochondria. By sequential addition of one or two of those inhibitors, nutrient dependencies and capacities of the corresponding pathways can be quantified. OCR (**J**) and ECAR (**L**) kinetic curves of 20 wo WT and *Taz*-KD pancreatic islets using sequential addition of BPTES (left), etomoxir (middle) or UK5099 (right). Starting with the addition of one of the inhibitors, followed by the addition of the remaining two to observe the nutrient dependency of the inhibited pathway. OCR (**K**) and ECAR (**M**) kinetic curves of 20 wo WT and *Taz*-KD pancreatic islets using sequential addition of UK5099 and etomoxir (left), UK5099 and BTES (middle) or etomoxir and BPTES (right). Starting with the addition of two of the inhibitors, followed by the addition of the remaining one to observe the nutrient capacity of the non-inhibited pathway. n (WT) = 10, n (*Taz*-KD) = 6, n number of experiments include N = 4 (number of animals). Data represent mean ± SEM (indicated by error bars); n and N numbers indicate number of experiments and animals; statistical significance was determined by unpaired Student *t* test: *p < 0.05, **p < 0.01, ***p < 0.001. Abbreviations: weeks of age (wo), *Tafazzin*-Knockdown (*Taz*-KD), Wildtype (WT), oxygen consumption rate (OCR), extracellular acidification rate (ECAR), oligomycin (Oligo), antimycin A (Ant A), rotenone (Rot).

**Supplementary Figure 3:**
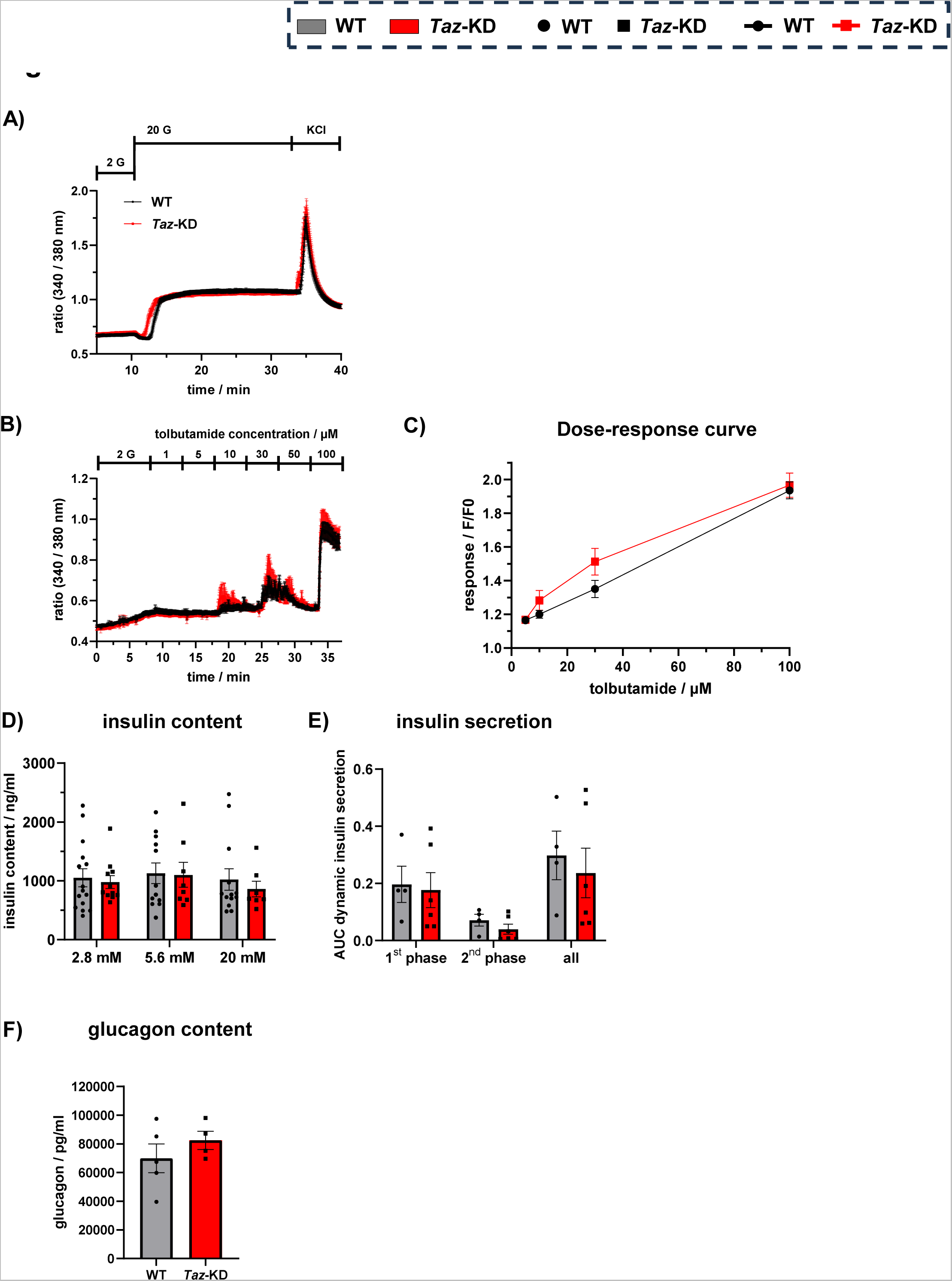
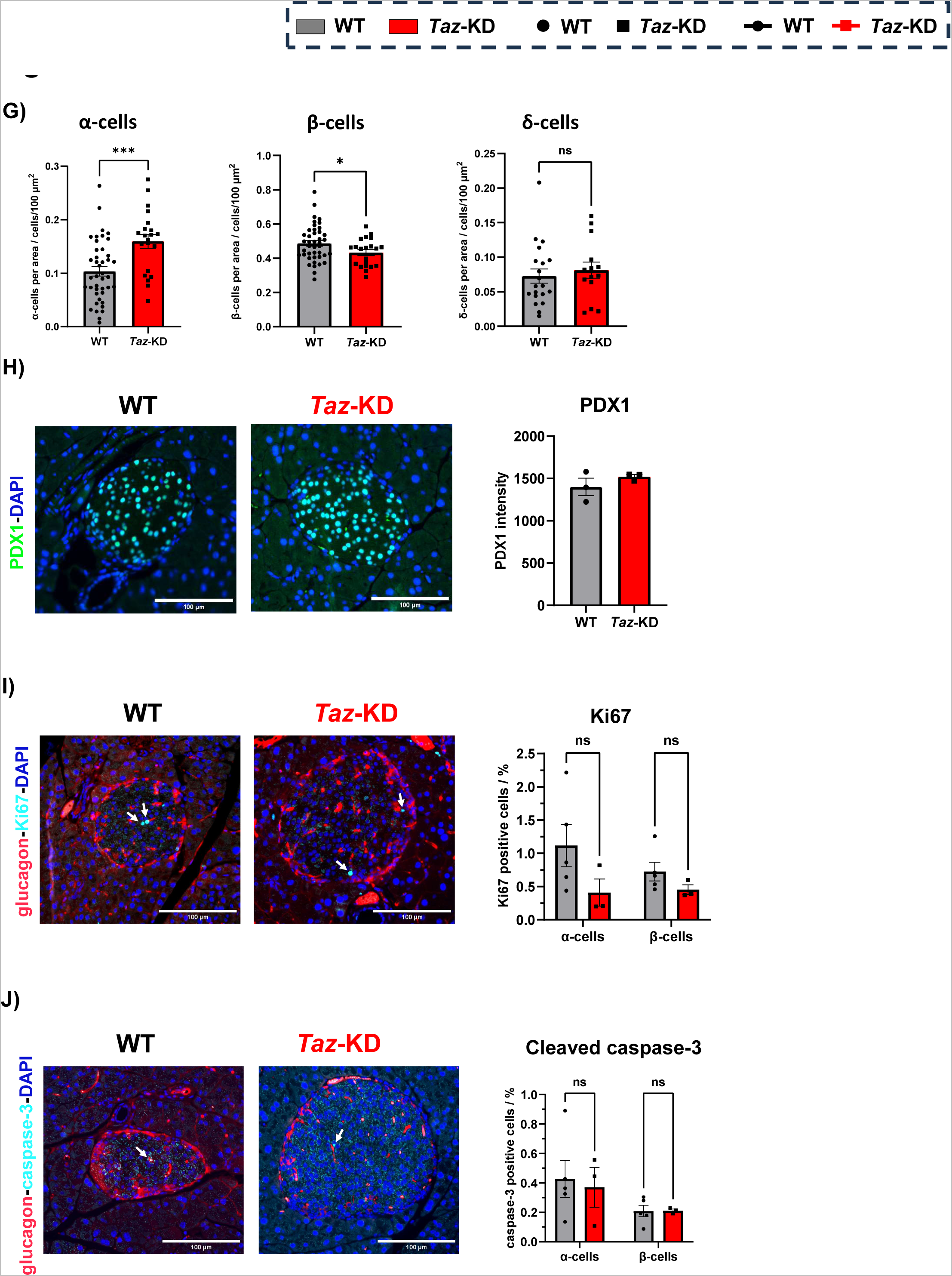
(**A**) Cytosolic calcium measurement of 20 wo WT and *Taz*-KD pancreatic islets using Fura-2 AM. The experiment represents the full-time course of the calcium experiment shown in Fig. 3B. (**B**) Cytosolic calcium levels of 20 wo WT and *Taz*-KD pancreatic islets with 2 mM glucose and increasing levels of tolbutamide (1 – 100 µM), N (WT) = 4, N (*Taz*-KD) = 6. (**C**) Calculated dose-response curve of tolbutamide addition and its resulting change in Fura-2 AM ratio (F) compared to the initial Fura-2 AM ratio (F0). n (WT) = 19, n (*Taz*-KD) = 22. (**D**) Quantification of insulin content of 20 wo WT and *Taz*-KD pancreatic islets at 2.8, 5.6 and 20 mM glucose concentrations, N (WT) = 16, N (*Taz*-KD) = 17. (**E**) AUC quantification of the dynamic GSIS separated in 1^st^ (10 – 20 min) and 2^nd^ (20 – 31 min) phase of insulin secretion, N (WT) = 4, N (*Taz*-KD) = 6. (**F**) Glucagon content of 20 wo WT and *Taz*-KD pancreatic islets, N (WT) = 5, N (*Taz*-KD) = 4. (**G**) Quantitative analysis of α- (left), β- (middle) and δ- (right) cell number of WT and *Taz*-KD pancreatic islets at 20 wo normalized to pancreatic islet area, n (α-cells, WT) = 42, n (α-cells, *Taz*-KD) = 21, n (β -cells, WT) = 42, n (β-cells, *Taz*-KD) = 21, n (δ-cells, WT) = 20, n (δ-cell, *Taz*-KD) = 14. Representative images and intensity quantification of IHC against PDX1 (**H**), Ki67 (**I**) and Caspase3 (**J**) in *Taz*-KD and WT pancreatic islets at 20 wo. Scale bar: 100 µm. N (PDX1) = 3, N (Ki67 and cleaved caspase-3, WT) = 5, N (Ki67 and cleaved caspase- 3, *Taz*-KD) = 3. Data represent mean ± SEM (indicated by error bars); n and N numbers indicate number of experiments and animals; statistical significance was determined by unpaired Student *t* test: *p < 0.05, **p < 0.01, ***p < 0.001. Abbreviations: weeks of age (wo), *Tafazzin*-Knockdown (*Taz*-KD), Wildtype (WT), oxygen consumption rate (OCR), extracellular acidification rate (ECAR), oligomycin (Oligo), basal rate (BR), immunohistochemistry (IHC), Pancreatic and duodenal homeobox 1 (PDX1), Glucose-stimulated-insulin-secretion (GSIS), area under the curve (AUC).

**Supplementary Figure 4:**
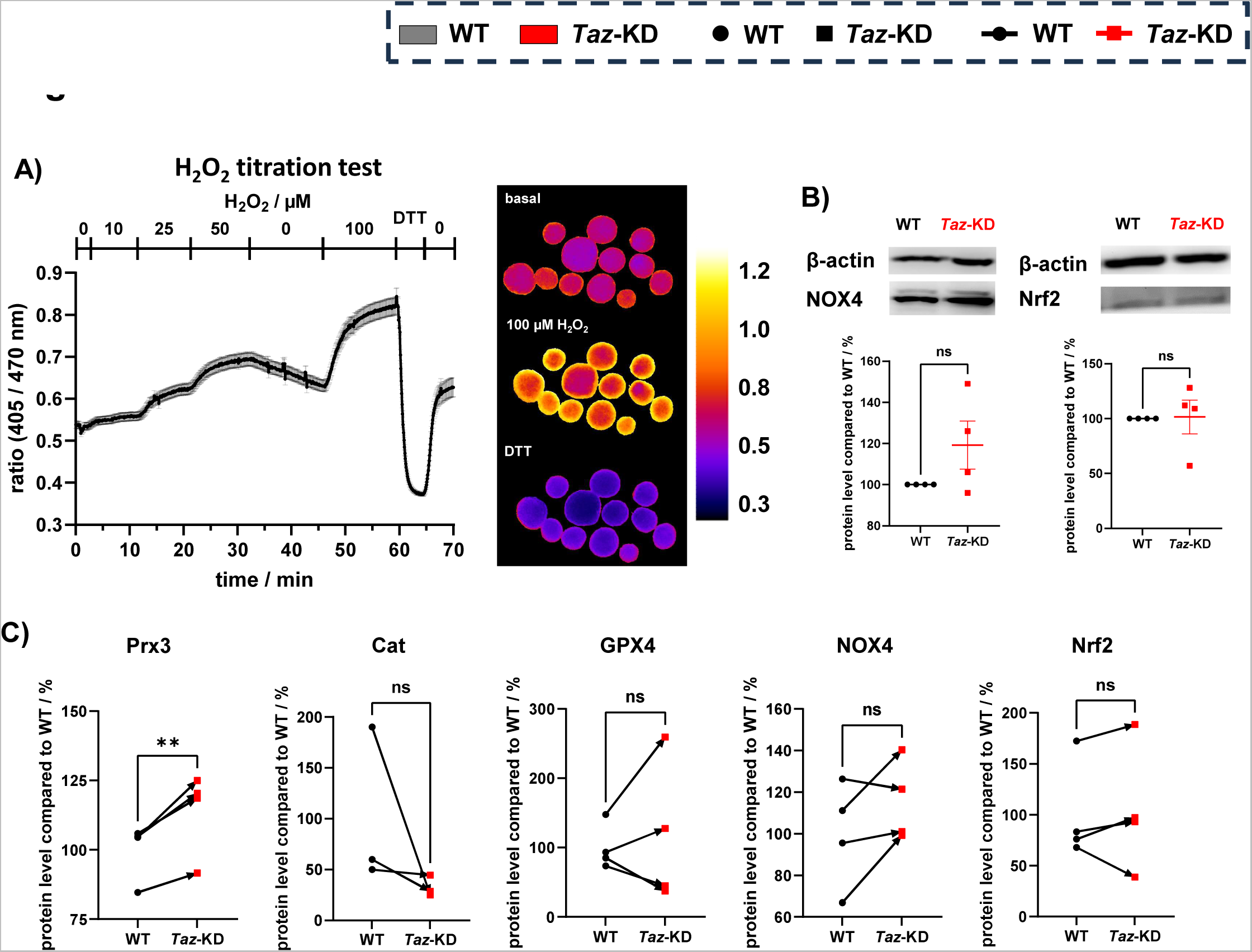
(A) Imaging of real-time H_2_O_2_ titration (0 – 100 µM) kinetics of isolated pancreatic islets heterozygous expressing of the mito-roGFP2-Orp1 sensor. Ratio (excitation: 405/470 nm, emission: 500 – 530 nm) images (right, Lookup table: “Fire”) created with ImageJ and reflect oxidation state at 0 µM (top), 100 µM H2O2 (middle) and 10 mM DTT (bottom). Calibration bar: redox state from 0.3 (reduced) to 1.2 (oxidized). n = 9 and N = 3. (**B**) Representative western blot and quantification of NOX4 (left) and Nrf2 (right) normalized to β-actin in pancreatic islets of 20 wo *Taz*-KD mice, N (NOX4) = 4, N (Nrf2) = 4. (**C**) Paired of western blot analysis of Prx3 (left), Cat (2^nd^ left), GPX4 (3^rd^ left), NOX4 (4^th^ left) and Nrf2 (right) normalized to β-actin in pancreatic islets of 20 wo *Taz*-KD mice, N (Prx3) = 4, N (Cat) = 3, N (GPX4) = 4, N (NOX4) = 4, N (Nrf2) = 4. Data represent mean ± SEM (indicated by error bars); n and N numbers indicate number of experiments and animals; statistical significance was determined by unpaired Student *t* test: **p < 0.01. Abbreviations: weeks of age (wo), *Tafazzin*-Knockdown (*Taz*-KD), Wildtype (WT), Dithiothreitol (DTT), area under the curve (AUC), catalase (Cat), NADPH oxidase 4 (NOX4), peroxiredoxin 3 (Prx3), glutathionperoxidase 4 (GPX4), GSK2795039 (GSK).

**Supplementary Figure 5:**
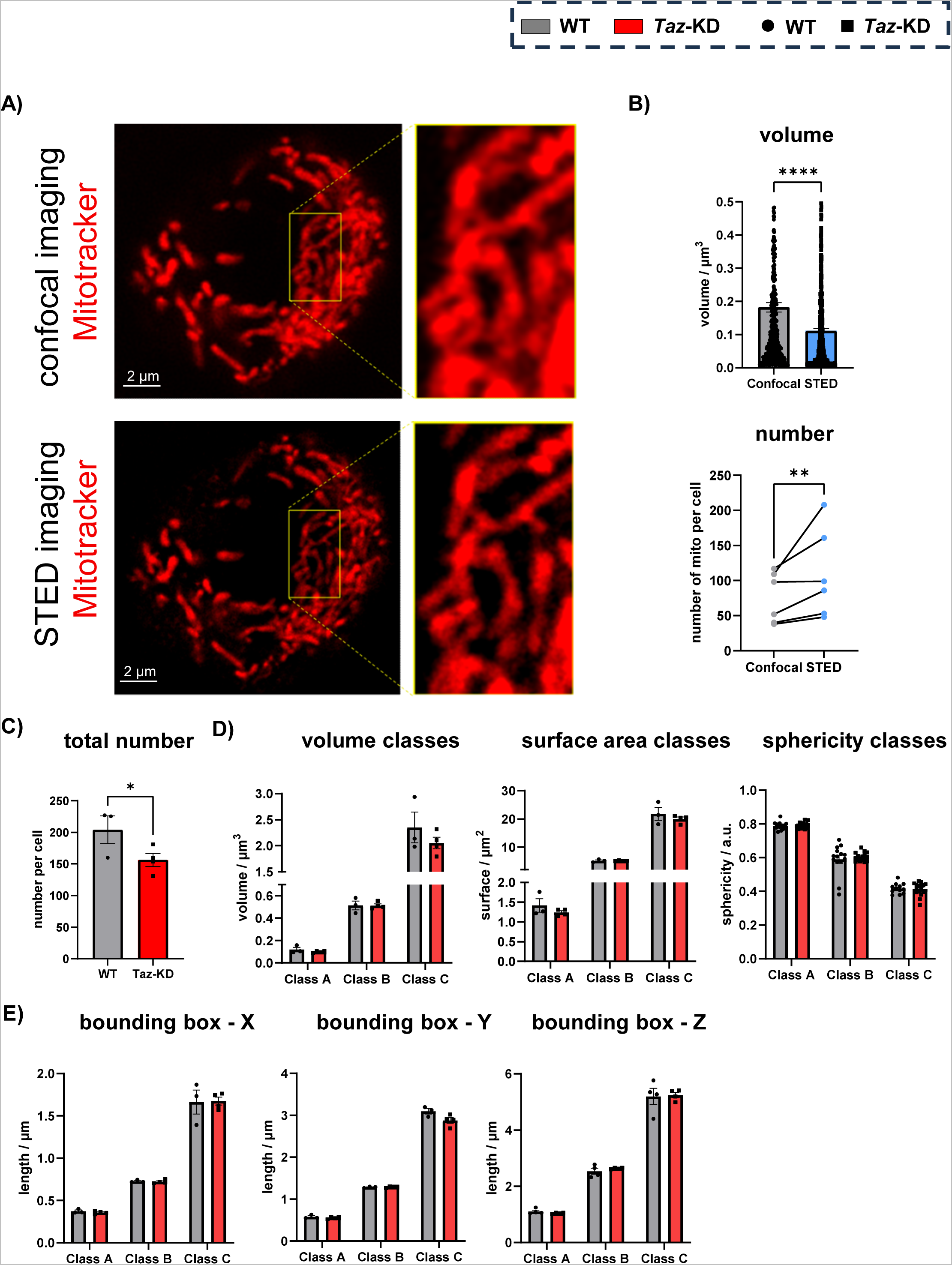
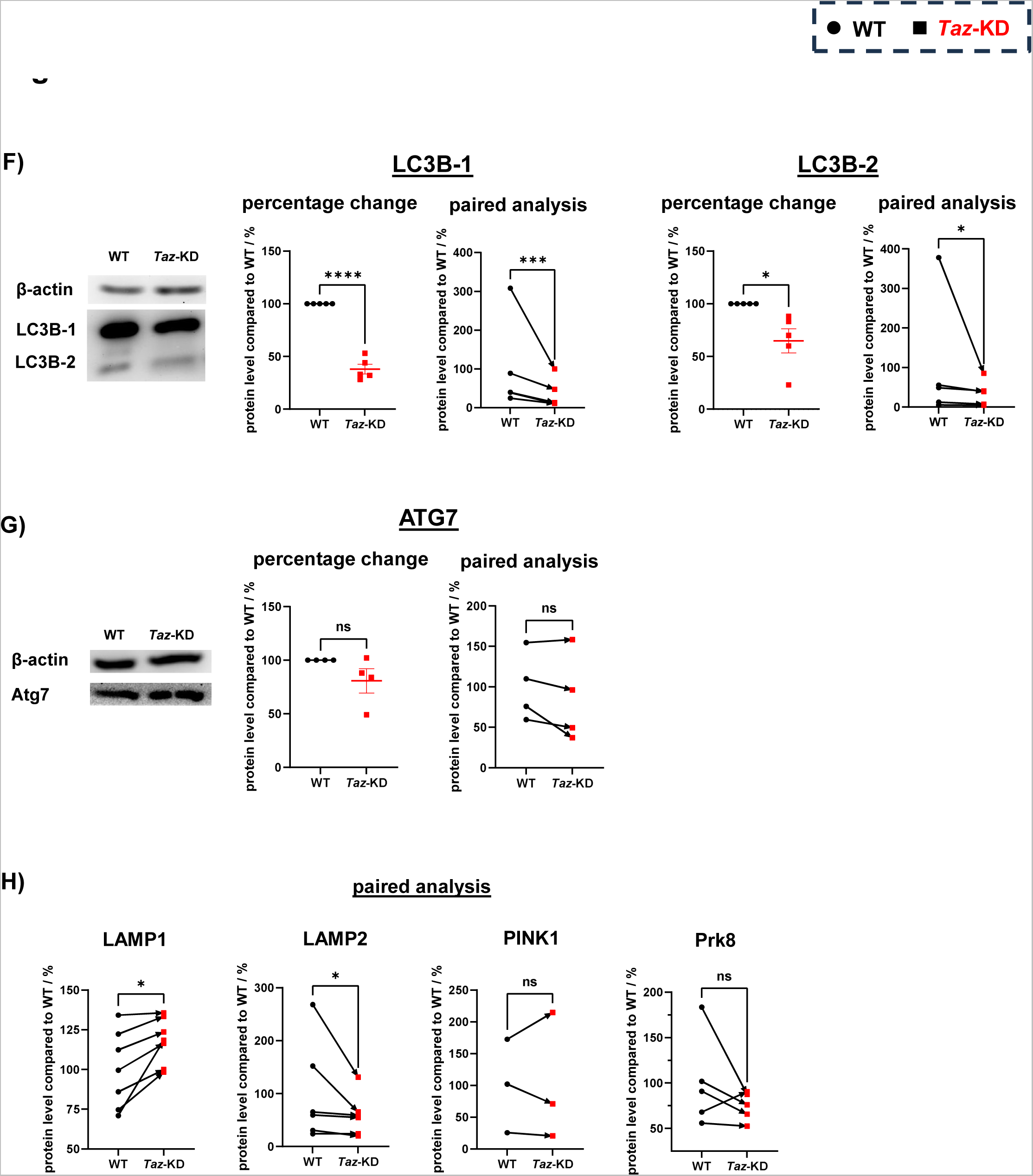
(**A**) Representative images of confocal (top left) and STED (bottom left) microscopy of mitochondrial network of the same pancreatic islet cell. Scale bar: 2 µm. (**B**) Comparison of single mitochondrial volume (top) and number (bottom) of confocal (black) and STED (blue) imaging. n = 6. (**C**) Total mitochondrial number per pancreatic islet cells from 20 wo WT and *Taz*-KD mice. N (WT) = 3, N (*Taz*-KD) = 4. Single mitochondrion (**D**) and bounding box (**E**) analysis separated by three surface area classes (A = 0.3 – 3 µm^2^, B = 3 – 10 µm^2^, C > 10 µm^2^) of 20 wo WT and *Taz*-KD dispersed pancreatic islet cells. The single mitochondria analysis includes the parameters volume (left), surface area (middle), sphericity (right) and the bounding box analysis includes the length in the three dimensions X (left), Y (middle), Z (right), N (WT) = 3, N (*Taz*-KD) = 4. Representative western blot, calculated percentage change and paired western blot analysis of LC3B-1 (left) and LC3B-2 (right) (**F**) and Atg7 (**G**) protein levels from 20 wo WT and *Taz*-KD pancreatic islets compared to β-actin protein level, N (LC3B) = 5, N (Atg7) = 4. (**H**) Paired western blot analysis of LAMP1 (left), LAMP2 (2^nd^ left), PINK1 (3^rd^ left) and Prk8 (right) normalized to β-actin in pancreatic islets of 20 wo *Taz*-KD mice, N (LC3B-2) = 5, N (LAMP1) = 7, N (LAMP2) = 6, N (PINK1) = 3, N (Prk8) = 5. Data represent mean ± SEM (indicated by error bars); N numbers indicate number of animals; statistical significance was determined by unpaired or paired (western blot) Student *t* test: *p < 0.05, ***p < 0.001, ****p < 0.0001. Abbreviations: weeks of age (wo), *Tafazzin*-Knockdown (*Taz*-KD), Wildtype (WT), lysosomal-associated membrane protein 1 (LAMP1), lysosomal-associated membrane protein 2 (LAMP2), parkin (Prk8).

**Supplementary Figure 6:**
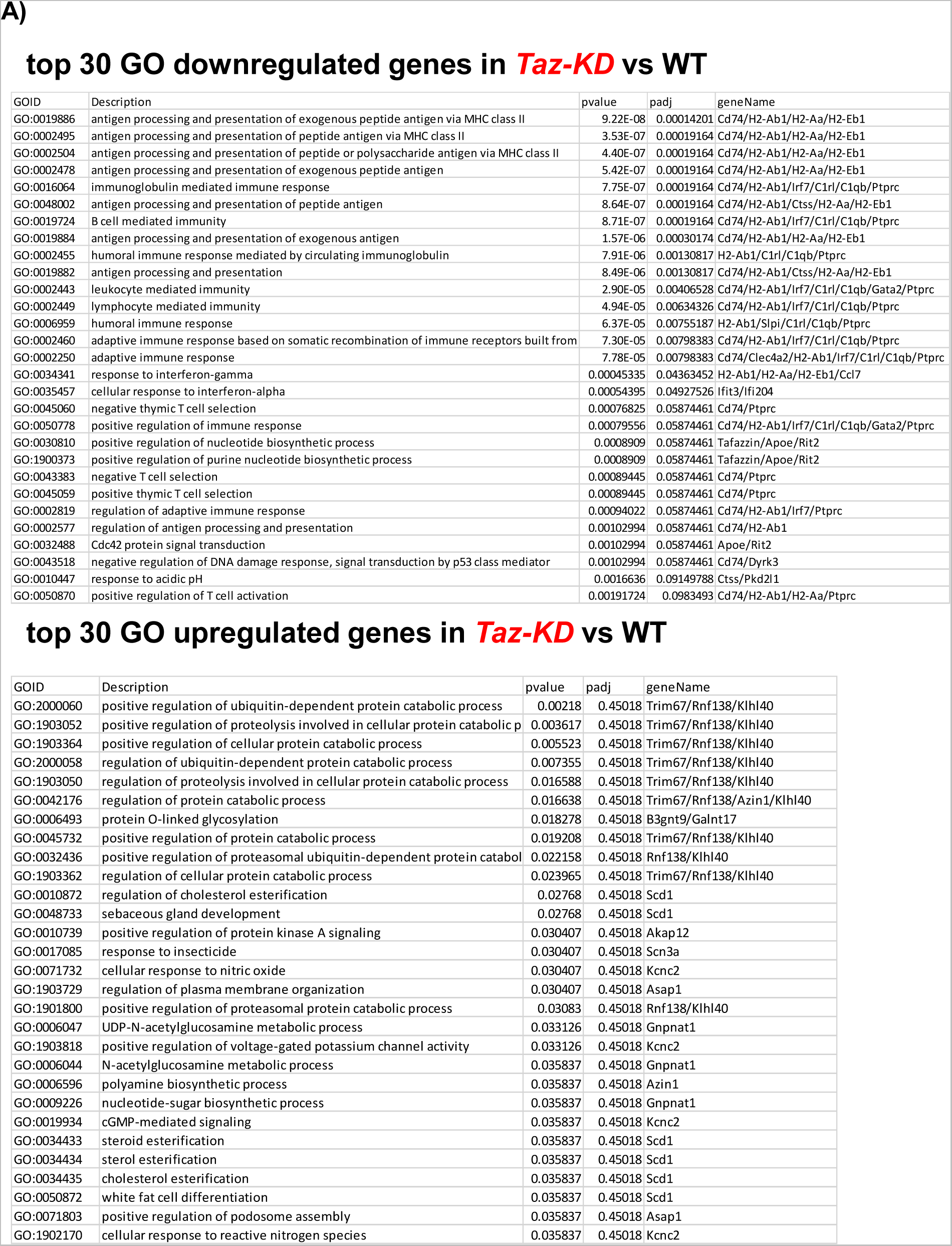
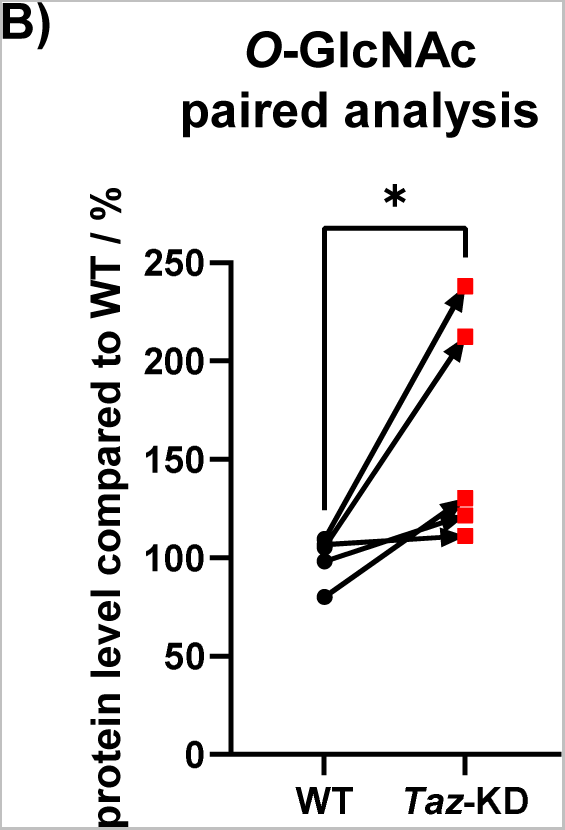
(A) 30 most significantly (selected by p values) downregulated (top) and upregulated (bottom) genes in GO pathway analysis of 20 wo *Taz*-KD pancreatic islets compared to WT controls. (B) Reactome analysis of enriched upregulated (left) and downregulated (right) genes of 20 wo *Taz*-KD pancreatic islets compared to WT controls. (**C**) Paired western blot analysis of *O*- GlcNAc normalized to β-actin in pancreatic islets of 20 wo *Taz*-KD mice, N = 5. Data represent mean ± SEM (indicated by error bars); N numbers indicate number of animals; statistical significance was determined by unpaired Student *t* test: *p < 0.05. Abbreviations: adjusted p-values (padj), gene ontology (GO), weeks of age (wo), *Tafazzin*-Knockdown (*Taz*-KD), Wildtype (WT), *O*-linked β-*N*-acetylglucosamine (*O*-GlcNAc).

**Supplementary Figure 7:**
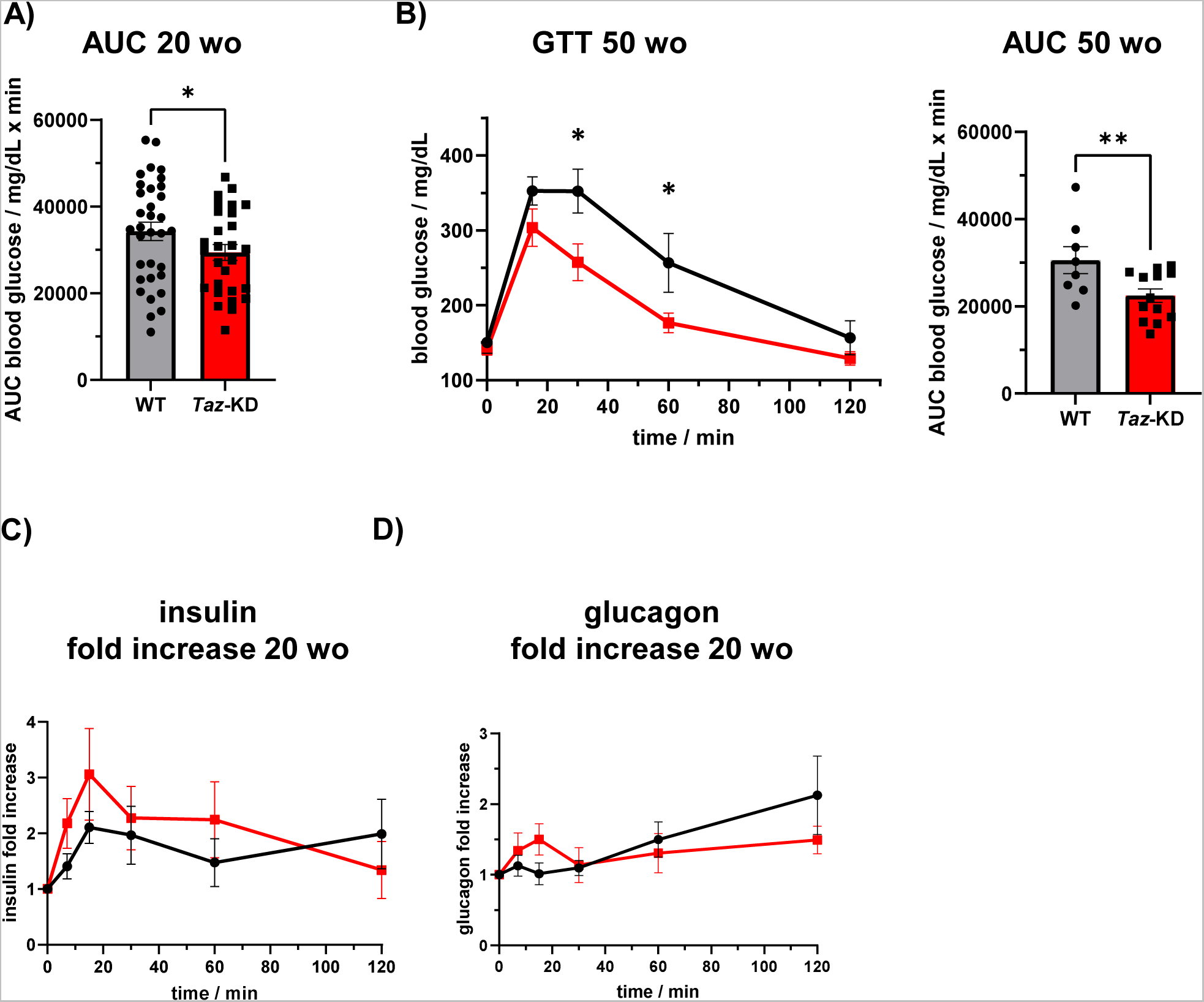
(**A**) AUC analysis of blood glucose levels of 20 wo WT and *Taz*-KD mice in GTT, N (20 wo, WT) = 24, N (20 wo, *Taz*-KD) = 22. (**B**) GTT analysis of blood glucose levels (left) and the quantified AUC (right) of 50 wo WT and *Taz*-KD mice, N (WT) = 8, N (*Taz*-KD) = 13. Calculated fold increase of plasma insulin (**C**) and glucagon (**D**) from GTT of 20 wo WT and *Taz*-KD mice, N (plasma insulin and glucagon, WT) = 7, N (plasma insulin and glucagon, *Taz*-KD) = 8. Data represent mean ± SEM (indicated by error bars); N numbers indicate number of animals; statistical significance was determined by unpaired Student *t* test: *p < 0.05, **p < 0.01. Abbreviations: weeks of age (wo), *Tafazzin*-Knockdown (*Taz*-KD), Wildtype (WT), glucose tolerance test (GTT), area under the curve (AUC).

## Notes

### Competing Interest Statement

Christoph Maack has received speaker honoraria and has been an advisor to Bristol Myers Squibb, Boehringer Ingelheim, AstraZeneca, Servier, Amgen, NovoNordisk, Bayer, Novartis, Edwards, and Berlin Chemie.
The other authors report that no conflict of interest exists.

